# Catalytic Mechanism and Differential Alarmone Regulation of a Conserved Stringent Nucleosidase

**DOI:** 10.1101/2025.04.13.648610

**Authors:** René L. Bærentsen, Kristina Kronborg, Ditlev E Brodersen, Yong E. Zhang

**Affiliations:** Department of Molecular Biology and Genetics, Aarhus University, DK-8000 Aarhus C, Denmark; Department of Biology, University of Copenhagen, DK-2200 Copenhagen, Denmark

**Author notes:** R. L. B. and K. K. contributed equally to this work.

**Keywords:** YgdH, PpnN, nucleosidase, LOG, cytokinin, alarmone, stringent response, ppGpp, pppGpp, persistence, cooperativity

## Abstract

Understanding how bacteria rapidly adapt their metabolism in response to external stimuli is key to addressing the present crisis of antibiotics-resistant infections. In the Gram-negative bacterium *Escherichia coli*, the universal stringent response is elicited in response to some antibiotics and involves production of the global alarmones, (p)ppGpp, which bind directly to many cellular targets. The nucleosidase PpnN that cleaves nucleotides into 5’-phosphate ribose and nucleobase, was shown to be a target of (p)ppGpp and control the delicate balance of bacterial fitness and persistence to fluoroquinolone antibiotics, thus conferring optimal survival strategies to bacteria during antibiotic selective pressure. Although both pppGpp and ppGpp stimulate the enzymatic activity of PpnN, they exert distinct effects on the enzyme’s cooperativity. The molecular mechanism underlying this subtle difference as well as the precise catalytic mechanism of PpnN, remain obscure. In this study, we provide mechanistic insights into the interaction of PpnN with substrate analogue, reaction products and alarmone molecules, which allows us to understand the catalytic mechanism of this family of nucleosidases and the differential modes of regulation by ppGpp and pppGpp, respectively. Comparison with the homologous LOG proteins involved in cytokinin production in plants reveals an ancient, universal mechanism for cleaving purine monophosphates that bacteria have incorporated regulatory controls through alarmones upon stringent responses.

**IMPORTANCE:** This study explores the distinct manners that the stringent alarmones pppGpp and ppGpp interact with the nucleosidase PpnN, which diverge from their typical, consistent effects on other target proteins. Furthermore, PpnN plays a key role in balancing bacterial fitness and tolerance to antibiotics, yet its catalytic mechanism has remained unclear. Through a combination of structural biology, molecular simulations, biochemistry, mutagenesis, and physiological analyses, this research uncovers the mechanistic differences in how both alarmones differentially impact enzyme cooperativity of PpnN and activation under stress conditions. Additionally, by analysing structures of PpnN complexed with substrate analogues, reaction products, and alarmones, as well as conducting bioinformatic comparisons, we propose a conserved catalytic mechanism shared with its homologue, LOG, a protein involved in cytokinin signalling - a critical growth hormone in plants. These insights underscore the nuanced regulatory roles of alarmones and PpnN/LOG group of nucleosidases, highlighting their evolutionary diversification to meet the varied environmental survival needs across organisms.

## INTRODUCTION

The universal stringent response, which is mediated by the alarmones guanosine penta- and tetra-phosphates (pppGpp and ppGpp, collectively referred to as (p)ppGpp), is triggered in bacteria in response to external stress such as nutrient deprivation and antibiotic exposure^1^. Once present at elevated levels, (p)ppGpp influences numerous vital physiological processes by interacting directly with specific target proteins^2,3^, leading to a change in transcriptional profile^4–6^, reduction in cell growth rate^7^, and an upregulation of stress response proteins. In addition to modulating these fundamental processes, (p)ppGpp plays crucial roles in bacterial virulence^8^, antibiotic tolerance and persistence^9^, highlighting the importance of the stringent response for understanding infectious diseases and tackling the antimicrobial resistance crisis. Consequently, gaining detailed insights into the molecular mechanisms of (p)ppGpp interaction and effects on target proteins hold great potential for developing novel antimicrobial treatment strategies in the future.

The nucleosidase PpnN (previously YgdH) is one of numerous novel target proteins of (p)ppGpp identified in the Gram-negative bacterium *Escherichia coli* (*E. coli*)^3^. PpnN is responsible for catalysing the cleavage of nucleotides into nucleobases and ribose 5’-phosphate (R5P)^10^ as part of the metabolic purine homeostasis, and thus functions in the opposite direction of the phosphoribosyl transferase enzymes, Hpt and Gpt, which are required for purine and pyrimidine nucleotide biosynthesis by the salvage pathway^11,12^. Both pppGpp and ppGpp allosterically stimulate the catalytic activity of tetrameric PpnN by binding at a dimer interface and inducing a large structural change that exposes the active site^13^. Simultaneously, (p)ppGpp inhibits the salvage pathway through direct interaction with Hpt, Gpt^3^, and the recently discovered (p)ppGpp-binding partners Gsk^14^ and PurF^2^. Stimulation of PpnN activity thus acts in synergy with the inhibition of the salvage (Hpt/Gpt/Gsk) and *de novo* (PurF) nucleotide biosynthesis pathways, leading to an overall reduced nucleotide pool and slower cell growth as well as redirection of resources^15^ towards stress responsive functions. Together, these observations support the idea that (p)ppGpp and PpnN participate in coordinating competitive fitness and antibiotic persistence in organisms such as *E. coli*^13^.

Although we have a good mechanistic and structural understanding of how *ppp*Gpp stimulates PpnN, two major questions remain to be answered: First, despite representing a large class of similar enzymes, the catalytic mechanism of PpnN is still not fully understood. Sequence similarity suggests a relationship with the distant group of Lonely Guy (LOG) enzymes in plants^16^, which produce cytokine-type hormones through cleavage of nucleotide-derived metabolites^17^. Therefore, a better understanding of the catalytic mechanism of PpnN is not only important for understanding the evolutionary connection between PpnN and the LOG proteins, but also the broader role of nucleosidase activity across different branches of life. Secondly, ppGpp and pppGpp have subtly distinct effects on PpnN activity. Apo-PpnN binds its substrate, GMP, cooperatively, as evidenced by a sigmoidal curve of enzyme kinetics^13^. Both ppGpp and pppGpp significantly stimulate the rate of the reaction; however, the presence of ppGpp abolishes cooperativity, while pppGpp significantly enhances the catalytic activity without affecting cooperativity^13^. It is worth noting that bacteria can produce varied ratios of pppGpp and ppGpp depending on the growth conditions^18–20^ and, both alarmones were shown to have differential potencies on affecting cell metabolism^21^. Further, in *E. coli* pppGpp can be readily converted to ppGpp via GppA^22^. Consequently, a detailed understanding of how the two molecules differentially affect enzyme function is important. While ppGpp and pppGpp often exert similar and additive effects on most target proteins, differential affinities have been observed for some target proteins, e.g., RF3^3,23^, EF4 (LepA)^3^, Der^3,24^, and RelQ^25^. Nevertheless, the distinctive effects of pppGpp and ppGpp on PpnN cooperativity are unique, and the underlying mechanism is not understood.

Here, we present new insights into the differential regulation of PpnN by both alarmones through determination of separate structures of PpnN in complex to pppGpp, ppGpp, guanine, and ribose 5-phosphate (R5P). Moreover, we analyse the structures of a catalytic mutant PpnN_E264A_ in complex with pppGpp and R5P; as well as wild type (wt) PpnN bound to ppGpp and 9-deaza-GMP (9dG), a GMP analogue. Using a combination of structural analysis and biochemical experiments, we identify the catalytic residues and propose a highly conserved catalytic mechanism for the PpnN and the LOG family of nucleosidases. Physiological experiments further confirm the critical role of PpnN catalytic activity in regulating competitive fitness and antibiotic persistence in *E. coli*. We further uncover a new allosteric binding site for 9dG, located at a separate subunit interface from the one that interacts with (p)ppGpp. Molecular dynamics simulations reveal that, in the presence of pppGpp but not of ppGpp, a specific loop region (loop_40-75_) changes its conformation and moves close to the dimer interface of PpnN where 9dG binds. In this process, new electrostatic interactions are established between loop_40-75_ and the 9dG-binding interface, which helps to stabilise loop_40-75_ in a new conformation, explaining the cooperative opening of the active sites in the presence of pppGpp, but not ppGpp. Together, these new data support molecular models for both the distinct roles of pppGpp and ppGpp, and the catalytic mechanism of a highly conserved family of nucleosidase.

## RESULTS

### Structure of alarmone-bound PpnN in a partially open state

Previous crystal structures of PpnN have either contained the enzyme tetramer in the *apo* (PDB 6GFL, *apo*-PpnN) or the fully activated form bound to four molecules of pppGpp (6GFM, 4:4 PpnN)^13^. Through continuous screening of PpnN incubated with pppGpp, we obtained a crystal form and determined the structure of PpnN in a partially allosteric state in which pppGpp was bound to three out of four PpnN molecules of the natural tetramer (3:4 PpnN, **Figure 1A** and **Table 1**). This conformation of the enzyme is overall similar to both *apo* and 4:4 PpnN; however, due to the partial binding of pppGpp, the tetramer is intrinsically asymmetrical and one monomer in the 3:4 PpnN structure shows a new, intermediate conformation. To better describe this, we define the uniform monomer conformation of *apo-*PpnN as the *closed* form, and the uniform monomer conformation of 4:4 PpnN as the *open* form (**Figure 1B**). Moreover, since pppGpp binds at a dimer interface, we will refer to the subunit that interacts with the 3’-end of pppGpp as its binding partner (**Figure 1C, s1a-c**). Using this, we can conclude that the three pppGpp binding monomers of the 3:4 PpnN structure are very similar to the 4:4 PpnN conformation, with an average root mean square deviation (rmsd) for Cα atoms of ∼0.4 Å compared to the open form. The corresponding value is 1.6-2.1 Å when compared to the closed form. The fourth monomer of the 3:4 PpnN structure is found in a partially open conformation (**Figure 1B**). The PAG1 domain (residue 1-150) of the partially open monomer more closely matches the open form (rmsd=0.35 Å) than the closed form (rmsd=1.37 Å); whereas the PAG2 domain (residue 330-454) more closely matches the closed conformation (rmsd = 0.67Å) compared to the open conformation (rmsd=1.76Å) (**Figure 1B**). Comparison of the pppGpp sites of the three monomers in the open form to 4:4 PpnN (**Figure 1C**) reveals that Glu69 in all cases is oriented towards the centre of the protein tetramer, which has previously been observed for pppGpp-bound PpnN^13^ (**Figure 1C**). Two of these alarmone sites have a clear density for pppGpp (**Figure s1a-b**), while the third open monomer that interacts with the 3’ phosphate group of a pppGpp, but not the 5’ group only has a weak density of pppGpp (**Figure s1c**). However, since Glu69 is clearly located towards the core of PpnN, we are confident in interpreting the site as being occupied by a pppGpp. Finally, the fourth alarmone site is in a conformation similar to *apo-P*pnN with Glu69 occupying the alarmone binding site (**Figure 1D, s1d**), consistent with no density for pppGpp (**Figure s1d**).

**Figure 1.**
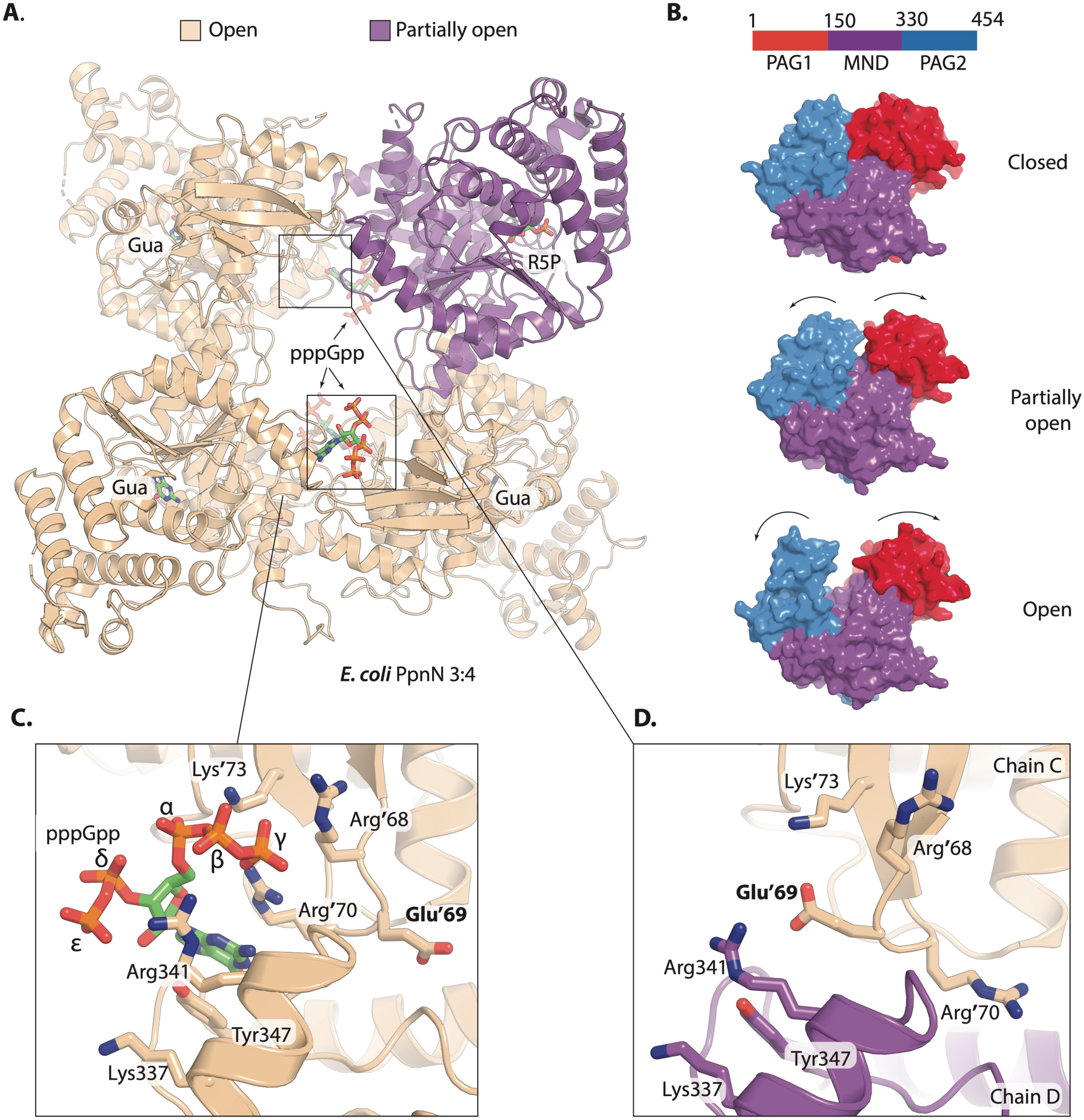
A new PpnN structure (3:4 PpnN) with a partially open conformation. **A)** Overview of the tetrameric PpnN. The individual subunits are coloured based on their conformations, with beige being the open conformation and violet being the partially open conformation. Gua, guanine; R5P, ribose-5-phosphate; pppGpp, guanosine pentaphosphate. All ligands are shown as stick-models. **B)** PpnN monomers in the closed, partially open and open conformations (PDB ID: 6GFL^13^ for the top panel, and PDB ID: 6GFM^13^ for the bottom panel), with the individual domains coloured separately (PAG1= red, monophosphate nucleosidase domain (MND) = purple, PAG2= blue). The thin black arrows indicate the movements of the PAG1 and PAG2 domains in the partially open and open conformations relevant to the closed conformation. **C)** pppGpp binds to the dimer interface as observed previously (PDB ID: 6GFM^13^), with the key coordinating residues shown in stick model and coloured. **D)** One of the four alarmone sites without pppGpp bound, with the key coordinating residues shown in stick model and coloured. Note the distinct positions of Glu69 and Arg70 in panel-**D** as compared to panel-**C**, without and with pppGpp bound respectively.

**Table 1.** Crystallographic data statistics.

|  | 3:4 PpnN (9HS1) | PpnN:ppGpp (9HS2) | PpnN E264A (9HS3) |
| --- | --- | --- | --- |
| **Data collection** |  |  |  |
| Wavelength (Å) | 0,97623 | 1,54180 | 0,97623 |
| Resolution range (Å) | 49,63-2,36<br>(2,45-2,36)* | 53,11-3,40<br>(3,52-3,40) | 47,42-3,5<br>(3,62-3,5) |
| Spacegroup | P4<sub>3</sub>2<sub>1</sub>2 | P4<sub>3</sub>2<sub>1</sub>2 | P4<sub>3</sub>2<sub>1</sub>2 |
| Unit cell dimensions |  |  |  |
| a, b, c (Å) | 152,66 152,66 224,54 | 155,12 155,12 226,09 | 156,92 156,92 224,65 |
| α, β, γ (°) | 90 90 90 | 90 90 90 | 90 90 90 |
| Total reflections | 217,448 (21,437) | 77,021 (7,557) | 72,038 (7,049) |
| Unique reflections | 108.735 (10.721) | 38.513 (3.779) | 36.032 (3.526) |
| Multiplicity | 2,0 (2,0) | 2,0 (2,0) | 2,0 (2,0) |
| R-meas | 0,06 (1,45) | 0,09 (1,64) | 0,18 (1,11) |
| Completeness (%) | 99,90 (99,95) | 99,35 (95,6) | 99,83 (99,6) |
| I/ΣSIGMAX(I) | 13,71 (0,66) | 11,11 (0,64) | 6,92 (1,01) |
| CC1/2 | 0,99 (0,36) | 0,99 (0,52) | 0,98 (0,52) |
| CC* | 1 (0,73) | 0,99 (0,83) | 0,99 (0,83) |
| **Refinement** |  |  |  |
| Reflections used in refinement | 108,656 (10675) | 38,301 (3,627) | 36,008 (3515) |
| Reflections used for R-free | 5438 (548) | 1916 (165) | 364 (36) |
| R-work | 0,19 (0,35) | 0,23 (0,42) | 0,22 (0,33) |
| R-free | 0,24 (0,38) | 0,29 (0,47) | 0,27 (0,38) |
| CC(work) | 0,97 (0,64) | 0,95 (0,67) | 0,95 (0,68) |
| CC(free) | 0,94 (0,53) | 0,92 (0,73) | 0,92 (0,49) |
| Number of non-hydrogen atoms | 14,475 | 14,063 | 14,254 |
| Protein (residues) | 1791 | 1765 | 1789 |
| RMS (bonds) | 0,007 | 0,007 | 0,0063 |
| RMS (angles) | 0,94 | 0,97 | 0,59 |
| Ramachandran statistics |  |  |  |
| Favoured (%) | 96,62 | 93,32 | 95,00 |
| Allowed (%) | 3,10 | 5,93 | 4,88 |
| Outliers (%) | 0,28 | 0,75 | 0,11 |
| Rama-Z score |  |  |  |
| whole | -0,65 (0,18) | -2,33 (0,19) | -1,48 (0,18) |
| helix | -0,31 (0,15) | -1,53 (0,14) | -0,71 (0,15) |
| sheet | -0,04 (0,31) | -1,53 (0,34) | -0,56 (0,33) |
| loop | -0,50 (0,50) | -1,65 (0,27) | -1,33 (0,25) |
| Rotamer outliers (%) | 2,24 | 0,00 | 0,07 |
| Clashscore | 8,55 | 11,81 | 13,83 |
| Average B factor | 67,10 | 156,53 | 104,88 |

\*Statistics for the highest-resolution shell are shown in parentheses

### PpnN has a highly conserved set of residues in the active site

PpnN was previously shown to be homologous to the group of Lonely Guy (LOG) proteins in plants, which function by converting the adenine-based plant precursors into active hormones through a nucleosidase reaction similar to the one carried out by PpnN^10^. LOG-like proteins have also been found outside the plant kingdom, including fungi and bacteria, i.e., suggesting it represents a more universal enzyme class^26,27^. PpnN belongs to PFAM PF03641, a family primarily consisting of putative lysine decarboxylases, encompassing over 60,000 proteins and including 22 experimental structures of 11 distinct proteins (**Figure s1e)**. These include PpnN full-length homologs from *Vibrio cholerae* and *Idomarina baltica* (PDB: 2PMB^28^ and 3BQ9^29^, respectively), a catalytic domain/LOG homolog from *Mycobacterium marinum* (PDB: 3SBX^30^), and homologous enzymes from fungi (*Claviceps purpurea*, PDB: 5AJU^30^) and plants (*Arabidopsis thaliana*, PDB: 2A33^31^). Several of these structures including *M. marinum* (3SBX^30^) and *P. aeruginosa* (5ZBK^32^) were determined with an AMP molecule inside their active sites (**Figure s2a**).

Inspection of the active sites of the 3:4 PpnN structure (**Figure 1A**) reveal additional electron density in all four subunits. The partially open PpnN monomer contains a bilobed density (**Figure s2b**), while the three fully open PpnN monomers have smaller, planar densities (**Figure s2c)**. Modelling of the products of PpnN cleavage of GMP, i.e., ribose-5-phosphate (R5P) into the bilobed density (**Figure 2A**), and guanine into the smaller densities (**Figure 2B**) led to satisfactory fits suggesting that cleavage of GMP took place in the crystal. Since no GMP was added in the crystallisation mixture, this could have been carried over with PpnN proteins purified from the expression strain. For the *P. aerugionosa* orthologue, a catalytic mechanism was proposed that involves stabilisation of the monophosphate nucleotide by two Thr residues (for 5’ phosphate and nucleobase), conserved Glu and Arg residues (for 2’ and 3’ OH) and activation of a water molecule by the conserved Glu of the conserved PGGxGTxx**<u>E</u>** motif^32^. Based on a structure based sequence analysis (**Figure s1e**) and the suite of enzyme:ligand structures (**Figure s1f-g, s2a**), we can predict that Glu223, Glu263, Glu264, Arg241, and potentially also His157 of PpnN might be critical for catalysis, although His157 is only conserved in PpnN-like proteins, but not LOG-like proteins.

**Figure 2.**
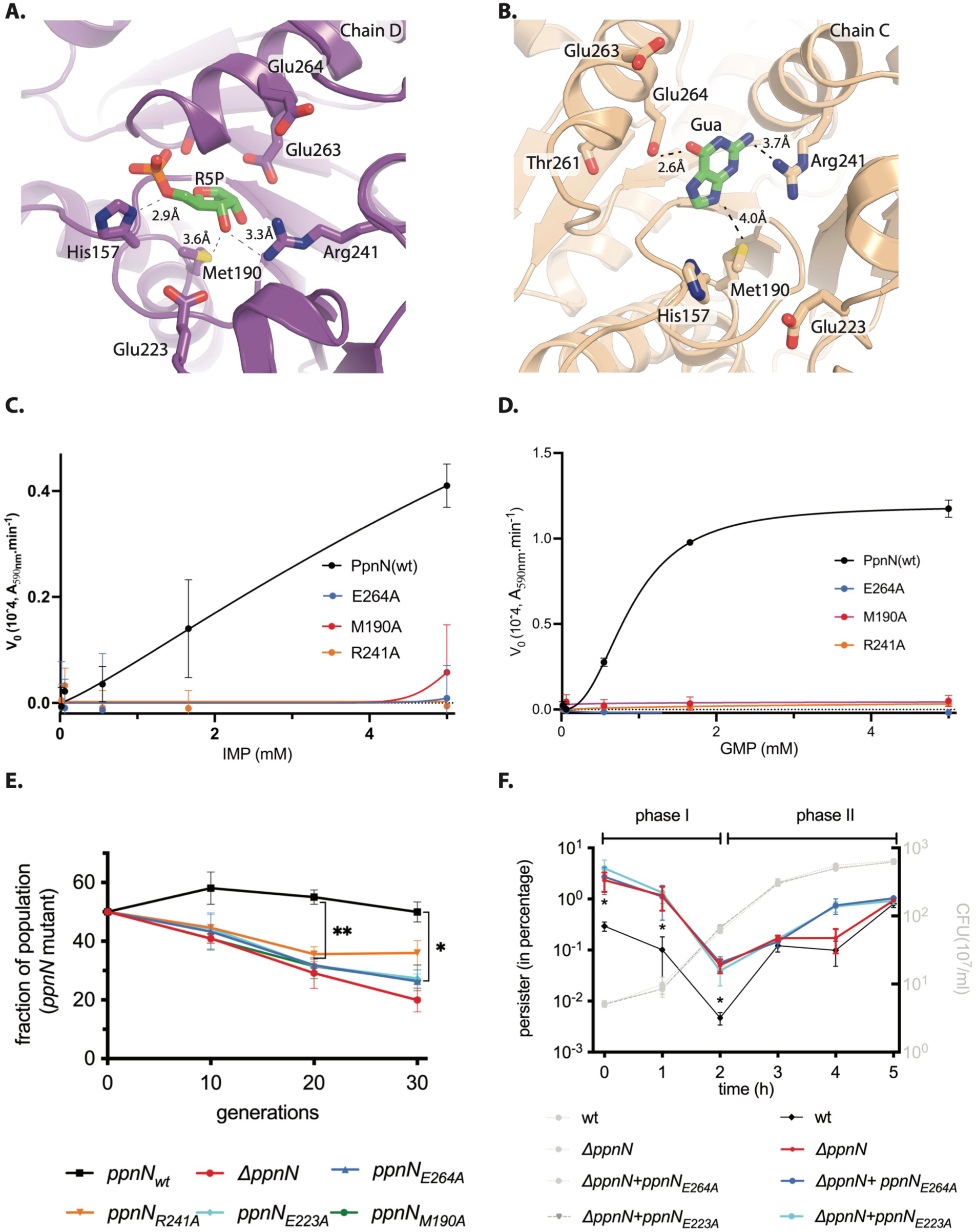
Catalytic activity of PpnN coordinates competitive growth and antibiotic tolerance. A, **B)** Active sites with the modelled products R5P and guanine reveal candidate residues critical for PpnN catalysis. **C, D)** Michaelis-Menton enzyme kinetic curves of wt and mutant PpnN proteins by using IMP (**C**) or GMP (**D**) as the substrate. Note the different scales of the Y-axes. At least three biological replicates were performed, and the curves are fitted using PRISM 10 and an allosteric sigmoidal model. **E)** Competitive one-to-one co-growth of various PpnN mutant strains with wild type MG1655 strain during feast-famine cycles (see Material and Methods for details). Three biological replicates were performed, and the average and SDs are plotted. *, **: student t-test p< 0.05 and <0.01, respectively. **F)** Persistence assay (left Y-axis) and growth curves (right Y-axis) of various PpnN mutant strains and the wild type MG1655 strain during regrowth from stationary phase cultures. Three biological replicates were performed, and the average and SDs are plotted. *, student t-test p< 0.05.

To test the roles of these residues in catalysis, we systematically changed each of them to alanine in individual PpnN protein constructs. To avoid contamination with endogenous *E. coli wt* PpnN, we expressed the constructs in an *E. coli* BL21 (DE3) Δ*ppnN* background before purification for biochemical analysis (**Figure s2d**). Initially, size exclusion chromatography (SEC) indicated that all variant PpnN proteins were soluble and behave like wt PpnN including forming a tetramer as expected (**Figure s2e**). We then performed kinetic assays with inosine monophosphate (IMP) as the substrate and followed the production of hypoxanthine, which can be quantified via an enzyme-coupled reaction as previously reported^33^. Using this approach, we found that the E264A, M190A, R241A and H157A mutations completely abolished catalytic activity, and no production of hypoxanthine could be detected even in the presence of 5 mM IMP (**Figure 2C** and **s2f**). The other mutations, i.e., E223A, T261A, E263A, appeared to be defective to various extents as compared to wt PpnN (**Figure s2f**). To corroborate these observations, we repeated the kinetic analysis with GMP as the substrate, as the product guanine can be quantitated via the same enzyme-coupled reaction as recently confirmed^34^ (**Figure 2D** and **s2g**). In this case, we obtain an EC_50_ of 0.89 ± 0.09 mM with a Hill coefficient of 2.5 ± 0.4, both values consistent with previous measurements by mass spectrometry (i.e., EC_50_ of 414 ± 14 !M, Hill coefficient of 2.8 ± 0.2)^13^, thus validating this method^34^. The E264A, R241A, E223A and M190A variants showed nearly no activity with GMP, while partial activities were observed for the E263A, H157A, T261A variants (**Figure 2D** and **s2g**). Altogether, these data confirm the structural and sequence analyses and suggest a highly conserved mechanism (see discussion below) between PpnN and LOG proteins.

### The catalytic activity of PpnN correlates with competitive fitness and antibiotic tolerance

It has previously been found that an *E. coli* Δ*ppnN* strain shows decreased fitness during feast-famine cycles of competitive growth against wt *E. coli* as well as increased tolerance to the fluoroquinolone antibiotics such as ofloxacin^13^. We therefore used the active site mutants of PpnN to ask if the catalytic activity *per se* is responsible for these two phenotypes. For this, we introduced the individual active site mutations into the endogenous *ppnN* allele of the *E. coli* MG1655 wt strain using the scarless method (see Methods for details)^35^. Furthermore, we inserted either a kanamycin (kan) or chloramphenicol (cam) resistance marker into the *attB* phage attachment site to allow for selection. To test fitness of PpnN catalytic mutants, we then grew the individual mutant strains with the *attB*::kan marker in competitive feast-famine cycles against wt *E. coli* MG1655 with an *attB*::cam marker and compared the fitness of the different strains in cycles of feast-famine^13^. To control this setup, we first confirmed that insertion of the kanamycin (or chloramphenicol) resistance marker into wt *E. coli* did not change its competitive fitness compared to the parental strain without the marker (**Figure 2E**, black line). We then found that the four mutant strains encoding the M190A, E223A, E264A, or R241A variants of PpnN showed significantly decreased fitness against wt cells, like the Δ*ppnN* deletion strain and thus consistent with their lack of catalytic activity (**Figure 2E)**.

A decreased competitive growth, however, might as well indicate an increased tolerance to antibiotics^13^. To test this, we measured the frequency of persister cells and sensitivity to ofloxacin for wt, Δ*ppnN*, and two mutant alleles, *ppnN_E223A_* and *ppnN_E264A_*, upon growth resumption from overnight stationary cells^13^. As reported before^13^, for all strains tested, we observed an initial drop of persisters (0-2 h, similarly defined as “phase I” in^13^) and their subsequent increases from 2-5 h (“phase II”, **Figure 2f**). This is expected since ofloxacin kills actively growing cells that have resuscitated from stationary non-growing state; while from 2 h, cells start entering the late-exponential phase, producing more persisters to ofloxacin. Importantly, during phase I, both *ppnN_E223A_* and *ppnN_E264A_* produced one order of magnitude higher persister cells relative to wt *E. coli,* indistinguishable from the Δ*ppnN* deletion strain (**Figure 2F)**. Together, these data show that the catalytic activity *per se* of PpnN correlates both competitive fitness and antibiotic tolerance in *E. coli*.

### A PpnN E264A structure reveals a partial conformation with product and pppGpp binding

We next attempted to capture a pre-cleavage state of PpnN by crystallising the inactive E264A variant in the presence of nucleotide substrates. Crystals formed in the presence of pppGpp and GMP (**Table 1**) diffracted to 3.5 Å and resulted in a well-refined structure (R/R_free_ = 0.22/0.27, referred to as E264A 4:4 PpnN), containing a complete PpnN tetramer in the crystallographic asymmetric unit similar to the 3:4 PpnN structure, suggesting asymmetry is present (**Figure 3A**). However, structural comparisons of individual subunits indicated that they adopt a partially open conformation, as reflected by the Cα rmsd values relative to other observed conformations: closed (0.74 Å), partially open (0.47 Å), and fully open (0.6 Å). In this structure, all four alarmone sites are occupied by pppGpp (**Figure s3a**) as in the 4:4 PpnN structure. Moreover, all four active sites show a similar bilobed density (**Figure s3b**) as observed in the 3:4 structure, likely corresponding to the product, R5P, and indicating that the partially open form represents a post-hydrolysis state (**Figure 3B**). The presence of R5P in the crystal structure of PpnN_E264A_ indicates that the E264A mutant has residual activity which was not observed during the timescale of the activity assay but happens during the time it takes for crystallisation to take place.

**Figure 3.**
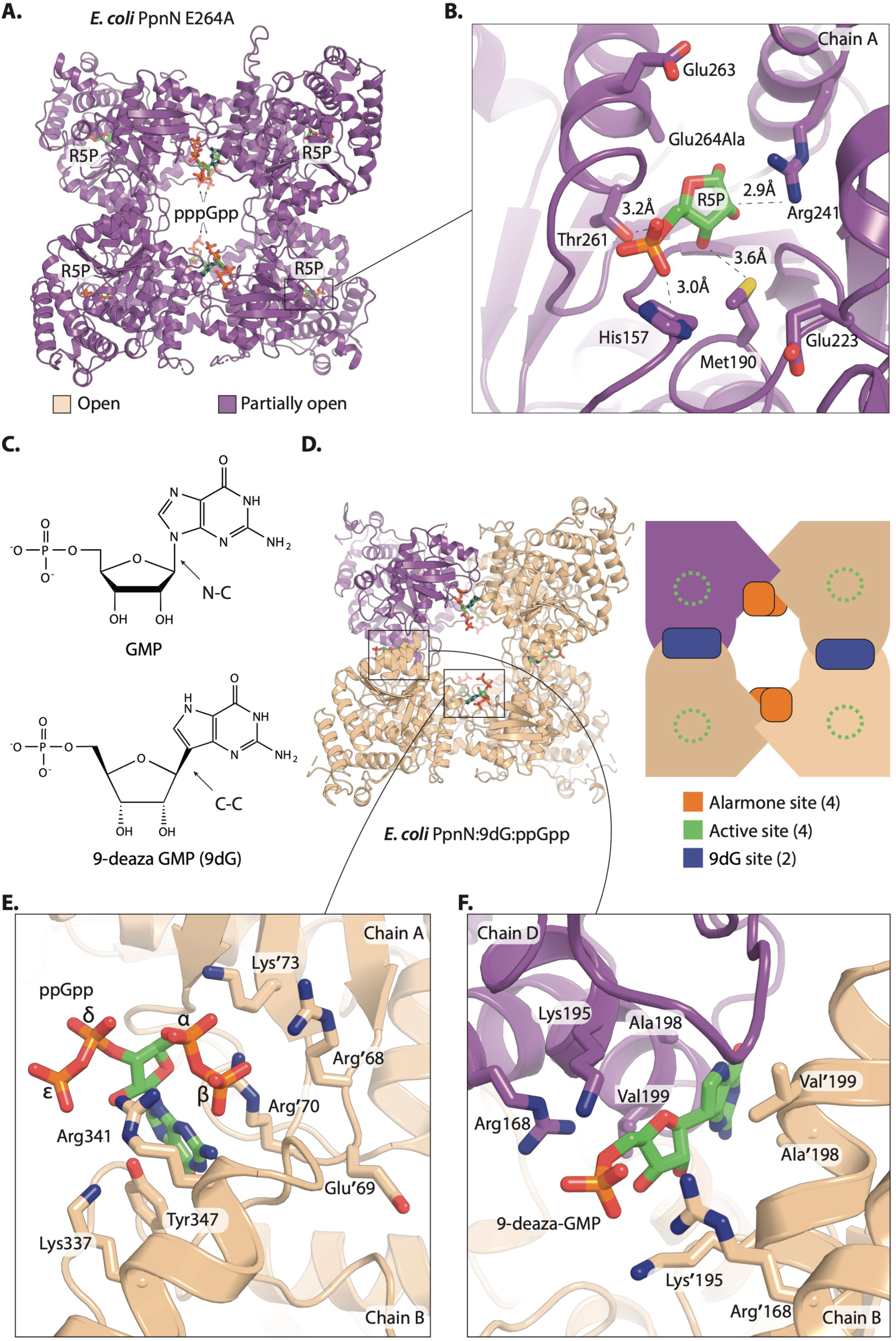
The first ppGpp-complexed PpnN structure reveals a new nucleotide binding site. **A)** Overview of the tetrameric PpnN_E264A_ structure with four pppGpp and four ribose-5’-phosphate (R5P) molecules bound. The purple colour indicate that all four monomers are in the partially open conformation. Ligands are shown in stick model. **B)** One representative PpnN active site from panel-**A** showing R5P and the coordinating residues (in stick model). **C)** Chemical structures of GMP and its analog 9-deaza GMP. Note the different C-C bond versus the N-C bond. **D)** The first PpnN structure in complex with ppGpp and 9-deaza-GMP (9dG). Beige monomers indicated open conformations whereas purple indicate a partially open conformation. (To the right side) A cartoon recap of the tetrameric PpnN containing four alarmone binding sites, four active sites and two nucleotide/9dG binding sites. **E)** One representative out of four ppGpp binding sites (from panel-**D**) to the same dimer interface as observed for pppGpp (Figure 1D), with the key coordinating residues shown in stick model and coloured. **F)** One 9dG binding site with modelled 9dG and coordinating residues shown in stick model.

### PpnN in complex with *pp*Gpp reveals an additional nucleotide binding site

To further attempt to capture PpnN in a pre-cleavage state, we co-crystallised PpnN_wt_ with 9-deaza-GMP (9dG), a GMP analogue containing a chemically stable glycosidic bond with a carbon atom at position 9 of guanine that likely prevents cleavage by PpnN (**Figure 3C**). Crystal screening with various combinations of analogues and alarmones resulted in a structure of PpnN_wt_ bound to 9dG and *pp*Gpp (3.4 Å, R/R_free_ = 0.23/0.27, **Table 1**, referred to as ppGpp-4:4 PpnN). This represents the first structure of PpnN in complex with *pp*Gpp. This crystal has a complete PpnN tetramer in the asymmetric unit (**Figure 3D**) and inspection of the structure shows that three subunits are in an open form and the remaining in a partially open form with Cα rmsd values of ∼0.5, ∼1.0, and ∼0.45 Å when comparing to the closed, open and partially open forms, respectively. All four canonical alarmone binding sites are occupied by ppGpp (**Figure 3E)**, but based on the density, the ligand appears more flexible than pppGpp (**Figure s3c**).

To understand the structural differences between ppGpp and pppGpp, we analysed the interacting residues and compared them using Ligplot (**Figure s3d-e**). First, this analysis reveals that the conformation of the alarmones varies between subunits of the same structure, but generally pppGpp is bound in a more consistent way than ppGpp. So, while similar residues are involved in the interactions (Arg68, Arg70, Lys73, Lys337, Arg341, Asp345, and Tyr347), ppGpp adopts several conformations and interacts with varying residues. This suggests a more dynamic, unsynchronized binding of ppGpp, consistent with its two-fold lower binding affinity to PpnN than pppGpp^13^. Inspection of the active sites of the ppGpp-4:4 PpnN structure reveals no density that can be modelled as either substrate or product. However, we observed a new density at the second dimer interface of the PpnN tetramer, i.e., the one that does not form the alarmone binding site (**Figure 3D, Figure s3f-g**). At this site, an elongated electron density was found into which 9dG could be modelled in contact with several conserved residues (Lys195, Arg168) that could serve to neutralise the charge of the 9dG phosphate group. Other residues (Val199, Ala198) contribute to forming a conserved, slightly hydrophobic pocket for the guanine moiety (**Figure 3F**). It thus appears like tetrameric PpnN contains four active sites, four alarmone binding sites, and two additional sites that support the binding to 9dG-like nucleotides (**Figure 3D**).

### A flexible loop controls the differential effects of both alarmones on PpnN cooperativity

Based on the structures of PpnN bound to ppGpp and pppGpp, we sought to understand how the two alarmones differentially regulate enzyme activity and cooperativity. To explore this, we used molecular dynamics to analyse whether there are differences in the flexibility and structural dynamics of the two alarmone-bound structures. For this purpose, we prepared both 3:4 PpnN and ppGpp-4:4 structures by removing 9dG, R5P and guanine prior to simulation, in order to focus on the effects of alarmone binding to PpnN. All simulations quickly stabilised (**Figure s4a-b**) and at the end of the simulations, all monomers in ppGpp-4:4 PpnN were in the fully open form, whereas the partially open monomer in 3:4 PpnN across all three simulations either opened immediately, after 200 ns, or stayed partially open for the duration of the simulation (**Figure s4c**). We first clustered the alarmone binding poses following the simulations. This gave a total of 31 binding poses for pppGpp and ppGpp, 18 shared and 13 unique to ppGpp. For the shared 18 binding poses, pppGpp predominantly existed in three; while ppGpp binds more dynamically and even escaped the binding site in one simulation (**Figure s4d-e, Table s4** for a definition of the observed binding modes). The observed variation in binding poses for the two alarmones thus correlates well with the weaker binding affinity of ppGpp^13^, the more diffuse electron density for ppGpp (**Figure s3c**), and the observed variations in ppGpp interactions with PpnN (**Figure s3d-e**).

Besides the alarmone binding site, the only other region showing significant structural changes during simulation is the loop covering residues 40-75 (loop_40-75_) of PpnN PAG1 domain (**Figure 4A)**. In the simulations of 3:4 PpnN, loop_40-75_ unwound and eventually stabilised in a new conformation in which Asp49 has established a new electrostatic interaction with Lys195 which is present in the central catalytic domain (residue 151-329) (**Figure 4B**). Importantly, loop_40-75_ forms half of the gate that restricts access to the active site in the closed form and moves away in the open form^13^ (**Figure s4f-g**). Although loop_40-75_ appears as a flexible region in all simulations, unwinding and subsequent stabilisation were only observed in 3:4 PpnN. Moreover, despite the 3:4 PpnN structure having overall higher resolution and low temperature factors, temperature factors are higher for loop_40-75_ suggesting that pppGpp induces more mobility of this loop (**Figure s4f-g**). Importantly, the new conformation of loop_40-75_ is incompatible with the binding of 9dG (**Figure 4C**). Altogether, this suggests that mobilisation and subsequent stabilisation of loop_40-75_ is promoted by binding of pppGpp, but not ppGpp.

**Figure 4.**
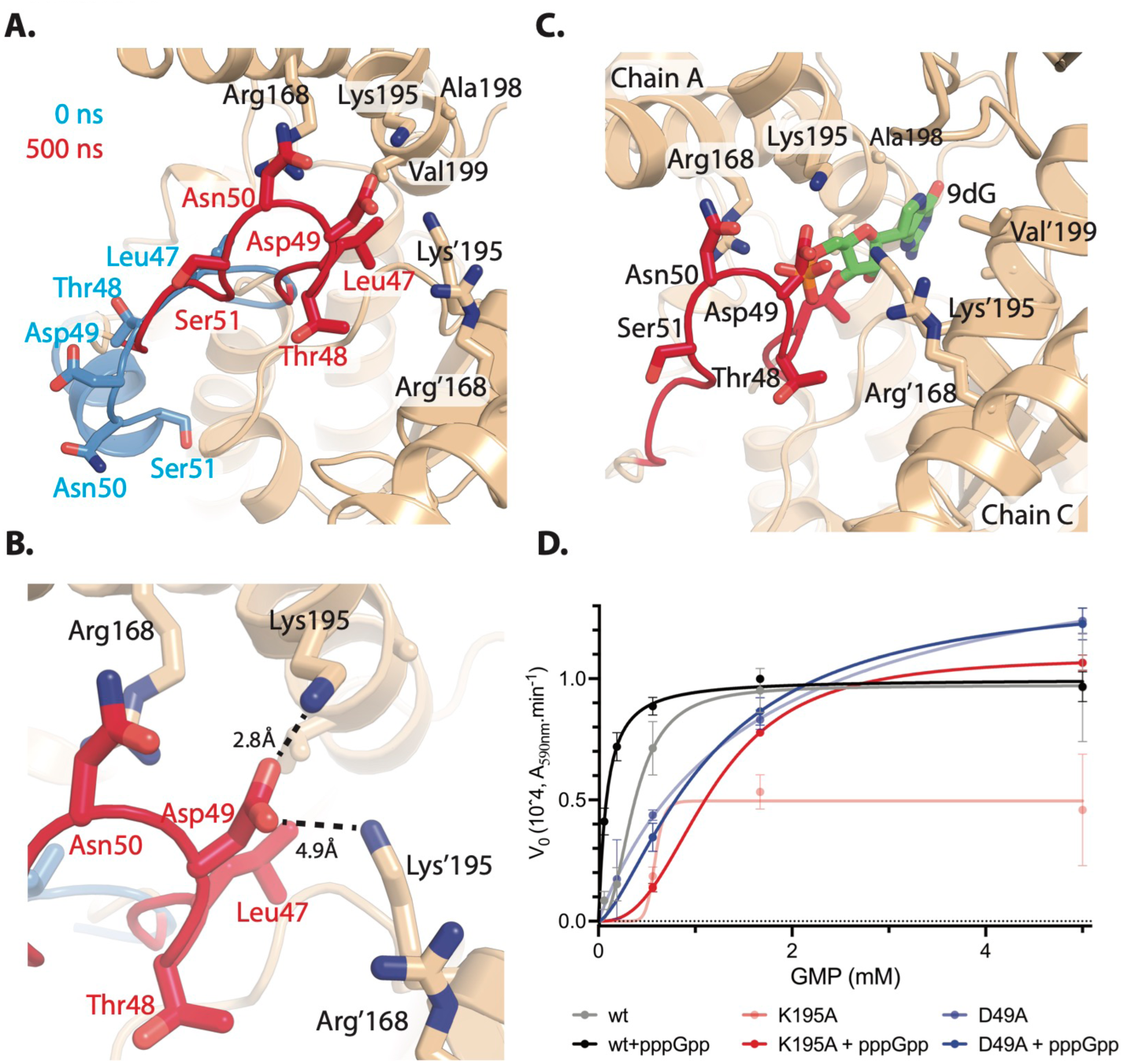
A dynamic loop underscores PpnN cooperativity without and with bound pppGpp. **A)** Conformations of residues 40-75 (loop_40-75_) before (blue) and after (red) molecular dynamic simulations (200 ns), upon only the binding of pppGpp. **B)** A new electrostatic interaction between Asp49 and Lys195 is established after simulation. **C)** Overlay of the alternative dimer interfaces with both the bound 9dG (ppGpp-4:4 structure, Figure 3D) and the stabilized loop_40-75_ (out of the simulation). Note the steric clash between 9dG and loop_40-75_. **D)** Michaelis-Menton enzyme kinetic curves of wt and mutant PpnN proteins in the presence (100 μM) or absence of pppGpp, using GMP as the substrate. At least three biological replicates were performed, and the curves are fitted using PRISM 10 and an allosteric sigmoidal model.

The above observed new and unique conformation of loop_40-75_ upon pppGpp binding might underlie its differential effect from ppGpp on maintaining PpnN enzyme cooperativity. To validate this, we changed Asp49 and Lys195 individually to alanine and studied enzyme activity and cooperativity in the presence or absence of pppGpp. Both variants (D49A and K195A) displayed similar SEC profiles as the wt PpnN suggesting that the tetramer remains intact (**Figure s4h-i**). Enzyme kinetics with GMP showed that PpnN_D49A_ has a hyperbolic, rather than a sigmoidal, Hill curve (Hill coefficient ∼ 0.98 versus 2.39 for PpnN_wt_) and a four-fold increased EC_50_ value (∼1.63 mM versus 0.37 mM for PpnN_wt_) (**Figure 4D**), supporting an important role of Asp49 in PpnN cooperativity and activity. Interestingly, pppGpp does not stimulate the catalytic activity of PpnN_D49A_. The EC_50_ value measured for PpnN_K195A_ in the absence of pppGpp increased almost two-fold to 0.58 mM as compared to PpnN_wt_. pppGpp-binding to PpnN_K195A_ resulted in a canonical sigmoidal curve (Hill coefficient ∼2.75), while the EC_50_ is ∼1.18 mM, i.e., 14 times that for PpnN_wt_ in the presence of pppGpp (0.084 mM). In conclusion, these data suggest that loop_40-75_ and, in particular, Asp49, plays a critical role for PpnN cooperativity in the presence and absence of pppGpp.

## DISCUSSION

### The dynamic behaviour of loop_40-75_ underscores the cooperative nature of PpnN

In this study, we present a series of PpnN structures, including the first structure of the enzyme in complex with ppGpp. Along with molecular simulations and biochemical assays, these structures allow us to elucidate mechanistic details about PpnN cooperativity and the differential response to the two alarmones, pppGpp and ppGpp. Based on these data, we propose a model (**Figure 5A**) where pppGpp binding to the alarmone site (1) induces a coordinated conformational change in loop_40-75_ (2), which contains the residues coordinating the 5’-end of (p)ppGpp, and Asp49. The conformational change promotes interaction between Asp49 and Lys195 (3), which opens and stabilizes the active sites, leading to cooperative activation of PpnN. In contrast, ppGpp disrupts enzyme cooperativity due to its more dynamic and uncoordinated binding mode, resulting from the absence of a 5’ "-phosphate.

**Figure 5.**
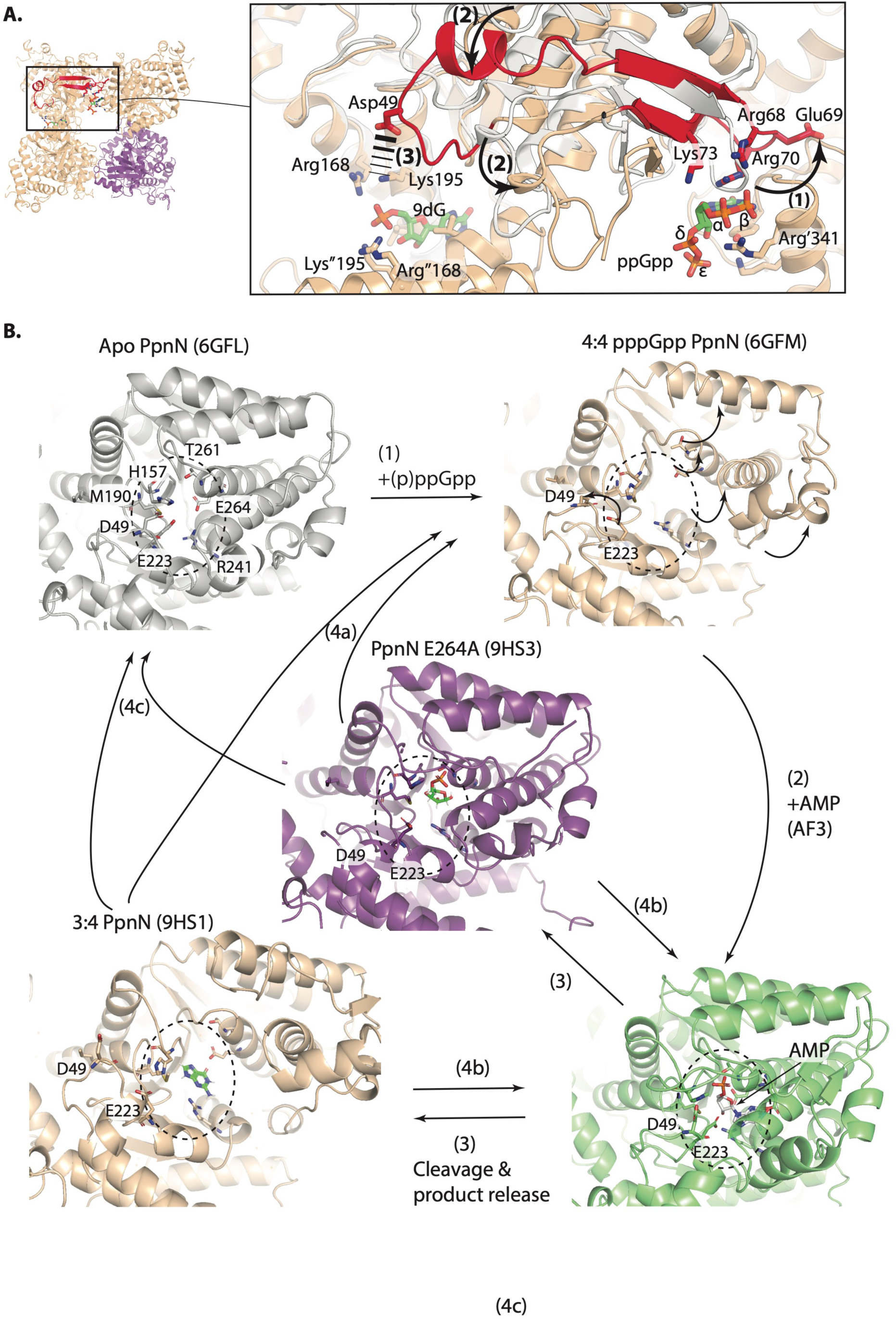
Proposed models of PpnN cooperativity and catalytic cycle. **A**) Model of differential alarmone activation of PpnN. For clarity, only a single site with ppGpp and 9dG (9-deaza-GMP) is shown, with the Apo form in grey and ppGpp bound form in beige. (1) The alarmone binds to the PpnN dimer interface, triggering a conformational change of the key coordinating residues (Arg68, Arg70, Arg341) and loop_40-75_ (highlighted in red) (2). Note that pppGpp induces a more pronounced conformational change compared to ppGpp (2). As a result, Asp49 is able to interact with the second dimer interface via an electrostatic interaction (3); whereas ppGpp, due to the lack of a 5’ "-phosphate, fails to induce this conformational change. **B**) Catalytic cycle of PpnN active site. The Apo state of PpnN (in grey) features a closed active site (marked by an oval with a dashed line). (1) Binding of alarmones activates PpnN (in beige) by moving away the PAG2 domain and loop_40-75_ to disclose and enlarge (indicated by arrows) the active site. (2) Nucleotides, modelled using AMP (in green) via Alphafold3, enter the active site. We predict that this binding results in the active site closing in to have the catalytic residues properly coordinating substrate for the cleavage reaction. Following cleavage, PpnN transitions (3) into either an R5P-bound partial open form (in purple) or a guanine-bound open form (in beige). PpnN can then adopt three possible states depending on alarmone binding and substrate availability: (4a) preparing for further substrate binding after product release; (4b) binding new substrates that outcompete retained products; (4c) returning to the Apo form due to alarmone dissociation and/or low substrate levels. Key residues studied in this work, along with substate and products, are shown in stick model and coloured. The active sites, marked by an oval, are fixed in size to illustrate the variations in active site dimensions across different states of PpnN. The corresponding PDB codes for each PpnN structure are indicated above the respective molecules.

Several pieces of evidence support this model. First, Ligplot (**Figure s3d-e**) revealed much more homogenous conformations of both pppGpp and its coordinating residues, than those of ppGpp. Consistently, more spurious electron density was observed for ppGpp, than for pppGpp (**Figure s3c**), and ppGpp binds to PpnN with a nearly two-times lower affinity than pppGpp^13^. Second, molecular simulation revealed three predominant binding modes of pppGpp, while many additional different modes were observed for ppGpp (**Figure s4d-e)**. Moreover, loop_40-75_ quickly mobilizes to the 9dG binding site upon binding of pppGpp, but not ppGpp (**Figure 4A-C)**. Consistently, loop_40-75_ has higher temperature factor in PpnN structures complexed with pppGpp than with ppGpp (**Figure s4f-g)**. Lastly, D49A mutation abolished the cooperativity of PpnN and its activation by pppGpp (**Figure 4D)**, suggesting the crucial role of D49 and loop_40-75_ in maintaining enzyme cooperativity by interacting with Lys195 at the 9dG binding site.

Altogether these data argue for a model where pppGpp binding activates PpnN by promoting a larger, more synchronous conformational shift of particularly the loop_40-75_; in contrast, ppGpp lacks a 5’ "-phosphate and appears to not promote this structural adjustment, resulting in loss of enzyme cooperativity. To our knowledge, this is the first time that distinct modes of actions of both alarmones are characterized for their target proteins. Instead, in all previously investigated target proteins including the four crystallized with both ppGpp and pppGpp^36–39^, ppGpp and pppGpp were shown to have similar and consistent effects on them, with, at the most, differential affinities and thus potencies.

### Catalytic mechanism and cycle of the PpnN (and LOG) family nucleosidases

Extensive comparison of the sequences and structures of PpnN and LOG proteins and biochemical studies in this study allow us to propose a highly conserved catalytic mechanism of the PpnN and LOG group of nucleosidases (**Figure s5a**). First, structures of PpnN (in complex with pppGpp and the reaction products R5P and/or guanine) and LOG proteins (in complex with various ligands (AMP, sulphate, phosphate, R5P)) showed that R241 and E223 (PpnN) and the counterpart residues R92 and E74 (in *P. aeruginosa* PAO1 PA4923, PaLOG, PDB 5ZBK) coordinate the 2’- and 3’-hydroxyl groups of substrates (e.g. GMP/AMP). Second, biochemical assays confirmed that E223/R241/E264 (PpnN) and the corresponding E74/R92/E115 (PaLOG)^32^ are essential for the catalytic activities of both PpnN and LOG, respectively. Lastly, protein sequence and structure comparisons showed that both these residues and their three-dimensional poses (**Figure s1e-g**) are highly conserved across PpnN and LOG homologs. The combined evidence suggests a shared catalytic mechanism as depicted in **Figure s5a.** Briefly, the highly conserved Arg241 and Glu223 of PpnN coordinates the 2’- and 3’-hydroxyl groups of substrate nucleotides; and, enzymatic catalysis proceeds with Glu264 activating a catalytic water molecule, which performs a nucleophilic attack on the anomeric carbon of the ribose ring, releasing nucleobase and R5P.

Besides a deeply conserved catalytic mechanism between PpnN and LOG proteins, we also see distinct regulatory features on enzyme activities that are reflected in their different domain architectures. Although unclear, LOG proteins appear to rely on domain dimerization to regulate enzyme activity^32^. For PpnN, several structures solved previously^13^ and in this study allow us to propose a relatively clear catalytic cycle (**Figure 5B**), accounting for how alarmone binding facilitates the catalytic reaction. In the Apo closed form (in grey), a tightly constrained active site (indicated by a broken-line circle) is maintained to limit substrate entry. Upon alarmone binding (step (1) in **Figure 5B**), a conformational change (of esp. loop_40-75_) is triggered (as in **Figure 5A**), enlarging the active site to permit substrate entry. Once substrate enters, the active site is constrained to reposition the catalytic residues back, coordinating and priming the substrate for the cleavage reaction (step (2) in **Figure 5B**). After cleavage of substrate (step (3) in **Figure 5B**), PpnN can transition to either the partially open form with R5P or the open form with guanine. Afterwards, PpnN may assume three different fates depending on the alarmone binding status and intracellular substrate concentrations: (4a) preparing for further substrate binding after product release; (4b) binding new substrates that outcompete the retained products; or (4c) returning to the Apo form due to the loss of alarmone binding and/or low substrate availability. Together, these steps illustrate how the activity of PpnN homologs is regulated by alarmone binding under various physiological conditions in the host bacteria.

### Outstanding questions

The new results reported here also raise several important questions. Would canonical nucleotides (e.g., GMP) bind to the same PpnN dimer interface that 9dG binds and regulate PpnN activity? If so, how would they influence PpnN activity and global cell metabolism under varying growth conditions in *E. coli*? Answering these questions will provide a deep understanding of *E. coli* physiology, since we have demonstrated that the catalytic activity *per se* of PpnN coordinates *E. coli’s* competitive growth and antibiotic tolerance (**Figure 2E,2F).** Furthermore, PpnN contributes to Salmonella’s tolerance to the host complement system^40^, and mutations in PpnN have been identified in patients with *E. coli*^41^ and Salmonella infections^42^. Additionally, an intriguing question is how the regulatory PAG1/2 domains evolved to regulate PpnN’s central nucleosidase activity in bacteria. Similarly, what mechanism regulates bacterial LOG-like proteins, which lack the PAG domains? Do LOG-like proteins respond to nucleotide-like molecules, such as (p)ppGpp, which plays a key role also within plastids? Further, LOG-like proteins appear to prefer larger, modified nucleotide substrates over the canonical nucleotides^43^. Could *E. coli* PpnN and its full-length homologs exhibit the opposite preference due to the steric hindrance of the PAG domains? Investigating these questions will deepen our understanding of bacterial nucleotide metabolism, their regulatory networks, and broader physiological roles in bacterial growth and pathogenicity^43^. Additionally, it could shed light on how (p)ppGpp and LOG proteins might interact and function together within plant plastids and contribute to overall plant physiology^17^.

### Footnotes

This study was supported by a Novo Nordisk Foundation Project Grant (NNF19OC0058331) and a Danmarks Frie Forskningsfond grant (2032-00030B) to Y.E.Z, and an Ascending Investigator (NNF18OC0030646) and a project grant (NNF17OC0028072) from the Novo Nordisk Foundation to D.E.B.

## ACKNOWLEDGEMENTS

We wish to thank Paulina Grucela for making the *E. coli* chromosomal mutant strains of *ppnN*. The authors are thankful to the beamline staff at Diamond Light Source for the *apo* structure of PpnN, and P13 at EMBL in Hamburg for the structure of pppGpp-bound PpnN and Sine Nøhr Nielsen for help with refinement of the pppGpp-bound structure.

## AUTHOR CONTRIBUTIONS

**a)** D. E. B and Y. Z. designed this study and acquired funding; R. L. B., K. K. and Y. Z. acquired experimental data; Y. Z., R. L. B. and D. E. B. wrote the draft manuscript; D. E. B. performed the bioinformatics analyses; and all authors analysed the data and edited the manuscript.

## STAR METHODS

### CONTACT FOR REAGENT AND RESOURCE SHARING

Further information and requests for resources and reagents should be directed to and will be fulfilled by Yong Zhang

### EXPERIMENTAL MODEL AND SUBJECT DETAILS

*E. coli* DH5α (Stratagene) was used for cloning, *E. coli* BL21 (DE3) (Novagen) for protein expression, and MG1655 for experiments requiring an *E. coli* wildtype strain background (see **Table S2** for further details). For cloning and protein expression for biochemistry, nutrient broth (Oxoid), with agar when appropriate, was used. The experiments of growth resumption, competitive fitness and persistence were performed in LB-B broth, containing 10 g tryptone (Oxoid), 5 g yeast extract (Oxoid), and 10 g NaCl per liter with pH adjusted to 7.43. When appropriate, chloramphenicol (25 µg/ml), ampicillin (100 µg/ml), and/or kanamycin (25 µg/ml) were used.

## METHOD DETAILS

### Plasmid constructions

To construct the plasmids pCA24N-*ppnN*(E263A), pCA24N-*ppnN*(E264A), pCA24N-*ppnN*(E263A E264A), pCA24N-*ppnN*(H157A), pCA24N-*ppnN*(M190A), pCA24N-*ppnN*(E223A), pCA24N-*ppnN*(R241A), pCA24N-*ppnN*(T261A), pCA24N-*ppnN*(K195A) and pCA24N-*ppnN*(D49A) quick-change mutagenesis was carried out using pCA24N-*ppnN* (wt) as template^13^ and the relevant primer pairs listed in **Table S3**. All plasmids were purified from *E. coli* DH5α and confirmed by sequencing (primers PYZ34/35, Eurofins genomics). For protein expression and purification, individual plasmids were transformed into *E. coli* BL21(DE3) *ppnN::kan* (see **Table S3** for details). The scarless method^35^ was applied to introduce the catalytically defective mutants to the endogenous *ppnN* allele as reported before^13^. Briefly, the strain YZ366 (see **Table S3** for details) was used which had the *ppnN* gene deleted and replaced with the SceI-Cam fragment. To introduce *ppnN* alleles containing the catalytic mutations, primer pairs pYZ451/PYZ213 and pYZ452/PYZ212 (for *ppnN(*M190A) allele), pYZ453/PYZ213 and pYZ454/PYZ212 (for *ppnN(*E223A) allele), pYZ456/PYZ213 and pYZ457/PYZ212 (for *ppnN(*R241A) allele), pYZ458/PYZ213 and pYZ459/PYZ212 (for *ppnN(*T261A) allele), were used to amplify and generate the mutant *ppnN* alleles to be electroporated into YZ366 to obtain *in situ* complemented strains. The *ppnN* sequences of the respective strains were confirmed by PCR amplification and sequencing.

### Protein purification for biochemical assays

*E. coli* Bl21 (DE3) strains (YZ785-YZ795 and YZ1699-1700) were purified as reported before^13^, with slight changes. Briefly, expression of N-terminally hexa-histidine tagged PpnN mutants was induced with 1 mM IPTG at early exponential growth. Pelleted cells were resuspended in lysis buffer (50 mM Tris-HCl pH 7.5, 150 mM NaCl, 10 mM imidazole, 5% glycerol, 5 mM beta-mercaptoethanol (BME)), lysed by sonication, and purified via Ni-NTA agarose resins (Qiagen). Bound PpnN was washed with wash buffer (50 mM Tris-HCl pH 7.5, 150 mM NaCl, 20 mM imidazole, 5% glycerol, 5 mM BME) and eluted with elution buffer (50 mM Tris-HCl pH 7.5, 150 mM NaCl, 500 mM imidazole, 5% glycerol, 5 mM BME.) Final separation was obtained using a Superdex 200 10/300 GL (GE Healthcare) column equilibrated in 50 mM Tris-HCl pH 7.5, 200 mM NaCl, 5% glycerol, 5 mM BME.

### Biochemical assays of PpnN activity and the effects of pppGpp

Catalytic activity of PpnN was determined via coupled enzyme reactions as described in^33,34^ using 150 nM PpnN and various concentrations of either IMP or GMP as the substrate. In summary, PpnN was first diluted into the reaction buffer (25 mM HEPES pH 7.5, 130 mM NaCl, 5 mM KCl, 1.5 mM MgCl_2_ and 0.3 μM BSA). Enzyme reaction was started by addition of substrate and terminated at selected timepoints by the addition of 10 μl 1 M Na_2_CO_3_. The pH of the reaction mixture was restored to neutral with 1 M HCl, and 20 µl of this sample was transferred to a Corning^TM^ 96-well solid black microplate with flat bottom (10022561, Fisher Scientific). To start the second and third reaction, 170 μl of the second reaction buffer (25 mM HEPES pH 7.5, 130 mM NaCl, 5 mM KCl, 1.5 mM CaCl_2_, 1 mM MgSO_4_, 5 mM glucose, 0.3 μM BSA, 1 U/ml HRP, 60 μM Amplex^TM^ Red Reagent and 0.15 U/ml xanthine oxidase (X2252-25UN, Sigma Aldrich)) was added to the samples and incubated for one hour at room temperature. Finally, fluorescence of the samples was quantified in a plate reader with an excitation of 545 nm and absorption of 590 nm. To assess the effect of *ppp*Gpp on PpnN activity, 100 μM *ppp*Gpp was included in the PpnN reaction buffer.

### Competitive growth assay

Similar as in^13^. Briefly, the OD_600nm_ of overnight cultures of each strain was measured and an equal OD_600nm_ of cells for each strain were mixed in appropriate combinations. The mixed cells were washed once with 1x Phosphate Buffered Saline (PBS) before inoculated into fresh LB-B broth with starting OD_600nm_ = 0.005. The initial fractions of each combined strain were determined by serial dilution of the mixed cells and plated on LB agar plates containing either kanamycin (25 μg/ml, for the mutant PpnN strains) or chloramphenicol (25 μg/ml, for the wt MG1655 strain). After every 24 hrs of co-growth at 37°C with agitation (160 rpm), the mixed cells were re-inoculated by 1/1000 dilution in fresh LB-B broth and the fractions of each strain in the populations were similarly determined.

### Persistence assay

Similar as in^13^. Briefly, an overnight culture of each strain was made in LB-B broth from which 120 μl of cells were inoculated into 12 mL fresh LB-B broth at room temperature. Cells were then grown in a water bath at 37°C with agitation (160 rpm). After every hour, 2 mL of cells were removed into lethal concentration of the antibiotic ofloxacin (5 μg/ml) and incubated at 37°C with agitation (160 rpm) for 5 hours. Meanwhile, the total number of colony-forming units (CFUs) before exposure to ofloxacin was determined by serial dilution, spotting, and incubation at 37°C for 24 h before counting. After 5 h of ofloxacin killing, 1.4 mL of cells were collected, washed once with 1.4 mL of PBS, and pelleted. The supernatant was removed completely after a final spin and the pelleted cells were resuspended in PBS, serial diluted and spotted on LB agar plate to measure the number of persisters, by determining the CFUs. For time zero samples, 2 mL of the inoculated cells were immediately removed to mix up with and killed by ofloxacin, and the total CFUs before and after ofloxacin killing were determined as above.

### Protein purification for crystallisation

*E. coli* BL21(DE3) *ppnN::kan* cells harbouring the plasmid of choice were grown in 2L LB media to an OD_600nm_ of 0.6 before induction with IPTG at 18°C overnight. Cells collected from 4L LB media were suspended in lysis buffer (50 mM Tris HCl pH 7.5, 300 mM NaCl, 10 mM imidazole, 5% glycerol, 5 mM BME) with an added 1 mM PMSF, and lysed via sonication, followed by centrifugation prior to being loaded onto a HisTrap FF Crude column (Cytiva). The column was washed in wash buffer (50 mM Tris HCl pH 7.5, 300 mM NaCl, 20 mM imidazole, 5 mM BME) before PpnN was eluted with elution buffer (50 mM Tris HCl pH 7.5, 300 mM NaCl, 200 mM imidazole, 5 mM BME). The HisTrap eluate underwent 6x dilution in buffer A (50 mM Tris HCl pH 9.0, 5 mM BME) prior to being loaded to a SourceQ column and eluted with an increasing gradient of buffer B (50 mM Tris HCl pH 9.0, 1M NaCl, 5 mM BME). The sample was concentrated to no more than 15 mg/ml before loading a maximum of 500 μL onto a Superdex 200 Increase 10/300 column (Cytiva) running with gel filtration buffer (20 mM Tris HCl pH 7.5, 100 mM NaCl, 5 mM BME) as a final purification step.

### Protein:ligand co-crystallization

The concentration of PpnN used for crystallisation was 6-8 mg/ml and 1-2 mM ligand was used for co-crystallisation trials. 1 mM 9-deaza-GMP was selected as a non-hydrolysable substrate analog to be crystallized with wt PpnN, whereas the active site mutant PpnN_E264A_ was co-crystallized with 2 mM GMP. The mixed protein:ligand solution was incubated for 1 hour at 4°C, before being centrifuged to pellet potential aggregate. Commercial screening was done with Swissci 2-well plates, with a reservoir of 70 μL, and a drop of either 0.2 μL buffer and 0.2 μL protein:ligand solution or a 0.1/0.1 μL mix. Plates were set up at both 4°C and 20°C. Crystals occurred after a week of incubation and stopped growing a couple of days later.

### Protein crystal data collection, processing and refinement

The dataset for wt PpnN with pppGpp was collected at EMBL Hamburg, Germany at the P14 beamline, while the datasets for wt PpnN with ppGpp and 9-deaza-GMP, and PpnN E264A with pppGpp and GMP were both collected at the MAXIV facility in Lund, Sweden, at the BioMAX beamline. Previous similar crystal morphology gave rise to a P4_3_2_1_2 dataset, and 1800 frames, with a 0.1° oscillation, were collected. Indexing, integration, and scaling were all performed using XDSGUI^44^ and the resolution cut-off was determined by balancing high completeness (>95%), stable I/σ (∼1.0), and a CC½ around 0.4. Spacegroup and asymmetric unit composition were verified using Pointless^45^ and Matthews_coef from the CCP4 suite^45^, and followed by phasing using molecular replacement using Phaser^46^. For PpnN:ppGpp:9-deaza-GMP, the open conformation of PpnN was most successful as a search model (PDB ID: 6GFM^13^), while in the case of PpnN E264A:pppGpp:GMP, the closed conformation (PDB ID: 6GFL^13^) was a better fit. Refinement was initially done in buster^47^, and after finishing building and adding water, the final refinement was done in phenix.refine^48^. Ligands not existing in the COOT^49^ library provided by REFMAC5^50^ were built either using the Simplified Molecular-Input line Entry System (SMILES) string as a starting platform, or in the case of 9-deaza-GMP, GMP was used as a starting point, and then edited manually using Lidia^51^ as implemented in Coot. Restraints for all ligands were generated using PRODRG^52^ from the CCP4 software suite, and manually inspected after one round of refinement, and edited, if necessary, to assure correct chemistry. To assist in the validity of placed ligand molecules polder maps were used to attain CC values for the placed ligands^53^. To validate ligand binding sites, we recalculated the density maps around the ligands with and without bulk solvent for all three structures and compared them using cross correlation using polder omit maps. This analysis indicates that all sites contain ligands (**Table S1**).

### Molecular dynamics system

Unless otherwise stated the simulation was done in GROMACS^54^ (v. 2019.4). The crystal structure of PpnN:pppGpp had no mutations but was built as the full-length protein by removing the N-terminal tag in the model, and building regions initially not modelled due to high flexibility guided and validated by the deposited structure of PpnN bound to pppGpp (PDB ID: 6GFM^13^). Protonation of PpnN was analysed using ProPKA^55^ (v. 3.1.0) comparing PpnN:pppGpp to the apo structure (PDB ID: 6GFL^13^). All chains had E264 protonated, which is also part of the PGGxGTxxE motif proposed to be part of the catalytic motif, albeit with lower pKa for the opened PpnN molecules. E264 in PpnN with a closed conformation was kept protonated during simulations. Residues border-lining protonation were E223, also part of the proposed catalytic site, and H157, proposed to coordinate to the 5’ phosphate of the substrate. The parameters for pppGpp were selected based on analogous molecules using the library from the CHARMM36 forcefield^56^ (July 2019). The guanine and ribose moiety were derived from guanine-triphosphate-gammasulphur, the 5’ triphosphate from ATP and the 3’ diphosphate from the 5’ diphosphates of ADP. The O3’ was parameterized as the O5’. pppGpp was placed in a box which was made by taking the maximal distances of pppGpp and adding 12 Å out to the border of the box. The box was filled with water following the TIP3P model. Na and Cl ions were used to neutralize the charges and put the system in a 0.1M concentration, to simulate the concentration of pppGpp binding^13^.

Similar parameterisation and system were built for ppGpp, with the only difference being that the 5’-diphosphate was derived from ADP. Energy minimization, NVT, NPT and a production MD of 500 ns were performed successfully for the ligand, showing Cl ions coating the phosphate arms, and the base was in the favoured anti conformation which is seen in almost all (p)ppGpp bound structures^57,58^. Bound to PpnN is pppGpp in the syn-conformation. A rotational scan performed in Maestro (2019, v.1) on the N-C bond between the ribose and the base on the lowest energy conformation from the production runs shows that there is a ∼1 kcal difference between the two states, but with a ∼5 kcal barrier in between. The fully parameterized pppGpp was built back into the PpnN system, which used the same principles for box size determination, water model, neutralization, and minimization. The output was split into three separate runs of NVT, NPT and production after minimizing the energy of the system.

### Molecular Dynamics analysis

Initial inspection of the trajectories in VMD^59^ showed that some regions changed conformation during the 500 ns run, and also that pppGpp and ppGpp were adopting multiple binding modes. RMSD plots as compared to the initial frame of the simulation were extracted using VMD, and the data was visualized using Prism. Using the simulations, each frame was aligned to the MND domain of the PpnN that bound (p)ppGpp with its PAG2 domain at t=0. An initial clustering where a new cluster was formed when the RMSD of (p)ppGpp exceeded 1.5 Å was made. This was followed by a subsequent internal clustering where the RMSD was minimized, and a merging of similar clusters was attempted. These grouped up clustered formed the basis of the dataset for PpnN binding modes to (p)ppGpp. The script was adapted to isolate the binding mode of each ligand binding site individually, the data collects information from 9 pppGpp binding sites, and 12 ppGpp binding sites. The high flexibility of both alarmones, especially in the 5’ and 3’ phosphate arms, led to several binding modes for each data collection. All binding modes present in under 1% of the simulation time were ignored, leaving 5-13 binding modes per individual binding site. These binding modes were inspected manually and grouped, since two binding modes that the script had recognized as different were similar upon inspection.

### Bioinformatics analysis and structure-based sequence alignment

Structural homologs of the central catalytic domain of PpnN, i.e. excluding the PAG1/2 domains involved in allosteric pppGpp binding were identified by extraction of all experimental structures in PFAM family PF03641 (Possible lysine decarboxylase). These comprise a total of 22 experimental structures covering 11 unique proteins. Sequence alignment was performed using Clustal Omega and formatted using SnapGene (Dotmatics) with additional manual annotation.

## QUANTIFICATION AND STATISTICAL ANALYSIS

For enzymatic kinetic assays of wild type and varied mutant PpnN proteins using IMP or GMP as substrate, at least three independent replicates were performed for each reaction. Linear regression was applied to obtain initial velocities with GraphPad Prism 10, and the Michalis-Menton curves are fitted with a non-linear allosteric sigmoidal model using Graphpad Prism 10. The student *t*-test was used for statistics when relevant.

For the competitive growth assay, different one-to-one combinations of experiments were performed in at least three independent replicates. In the persistence assay, the percentage of persisters was calculated by dividing the CFUs after antibiotic killing by the CFUs before killing. At least three independent replicates were performed for each strain and the student *t*-test was used for statistics.

## DATA AND SOFTWARE AVAILABILITY

The structures of PpnN deposited in the Protein Data Bank under ID codes 9HS1(*3:4 PpnN* form), 9HS2 (*ppGpp 4:4 PpnN* form) and 9HS3 (pppGpp-bound PpnN E264A form).

---

**Table S1.** Polder omit map analysis.

| Ligand with chain ID | CC(1,2) | CC(1,3) | CC(2,3) | Certainty of ligand |
| --- | --- | --- | --- | --- |
| **PpnN:pppGpp** |  |  |  |  |
| pppGpp (chain A) | **0.65(0.63)\*** | **0.88(0.85)** | **0.60(0.58)** | + |
| pppGpp (chain B) | **0.60(0.65)** | **0.94(0.90)** | **0.58(0.57)** | + |
| pppGpp (chain C) | **0.67(0.68)** | **0.81(0.80)** | **0.64(0.57)** | + |
| Guanine (chain A) | **0.85(0.89)** | **0.94(0.94)** | **0.84(0.87)** | + |
| Guanine (chain B) | **0.81(0.86)** | **0.95(0.93)** | **0.80(0.84)** | + |
| Guanine (chain C) | **0.87(0.92)** | **0.85(0.83)** | **0.83(0.85)** | + |
| Ribose-5-phosphate (chain D) | **0.75(0.79)** | **0.76(0.74)** | **0.73(0.73)** | (+) |
| **PpnN E264A** |  |  |  |  |
| pppGpp (chain A) | **0.42(0.33)** | **0.74(0.68)** | **0.18(0.12)** | (+) |
| pppGpp (chain B) | **0.60(0.58)** | **0.80(0.78)** | **0.54(0.52)** | + |
| pppGpp (chain C) | **0.56(0.54)** | **0.87(0.82)** | **0.49(0.44)** | + |
| pppGpp (chain D) | **0.64(0.57)** | **0.83(0.80)** | **0.53(0.46)** | + |
| Ribose-5-phosphate (chain A) | **0.70(0.77)** | **0.91(0.89)** | **0.65(0.68)** | + |
| Ribose-5-phosphate (chain B) | **0.69(0.73)** | **0.86(0.83)** | **0.64(0.64)** | + |
| Ribose-5-phosphate (chain C) | **0.70(0.77)** | **0.90(0.89)** | **0.63(0.66)** | + |
| Ribose-5-phosphate (chain D) | **0.64(0.69)** | **0.92(0.88)** | **0.52(0.54)** | + |
| **PpnN:ppGpp** |  |  |  |  |
| ppGpp (chain A) | **0.53(0.41)** | **0.84(0.83)** | **0.54(0.40)** | + |
| ppGpp (chain B) | **0.48(0.39)** | **0.77(0.75)** | **0.40(0.37)** | (+) |
| ppGpp (chain C) | **0.51(0.46)** | **0.90(0.83)** | **0.41(0.33)** | + |
| ppGpp (chain D) | **0.47(0.39)** | **0.78(0.75)** | **0.42(0.35)** | (+) |
| 9-deaza-GMP (chain A) | **0.60(0.62)** | **0.86(0.83)** | **0.52(0.50)** | + |
| 9-deaza-GMP (chain C) | **0.62(0.64)** | **0.89(0.88)** | **0.63(0.63)** | + |

**Table S2:**
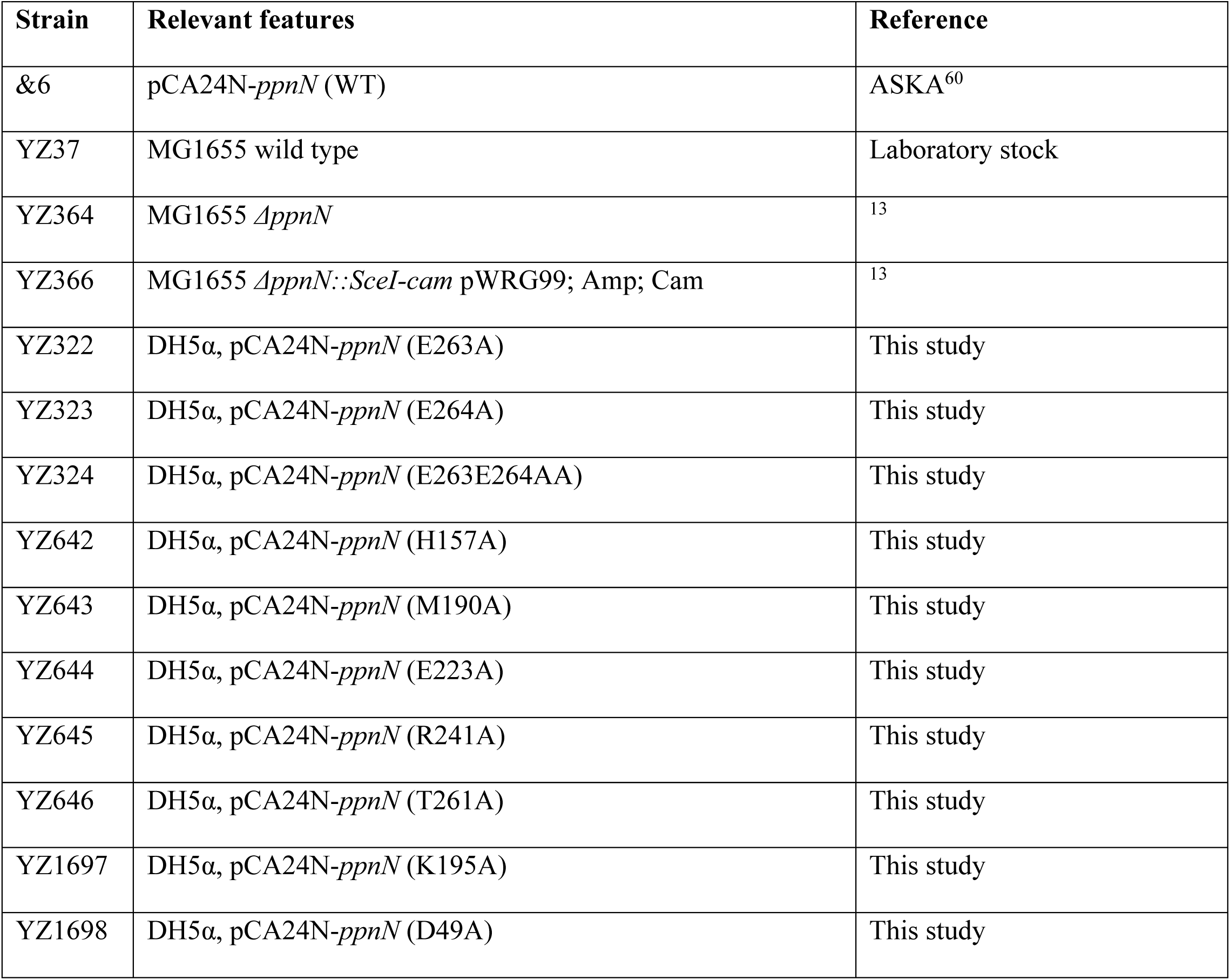
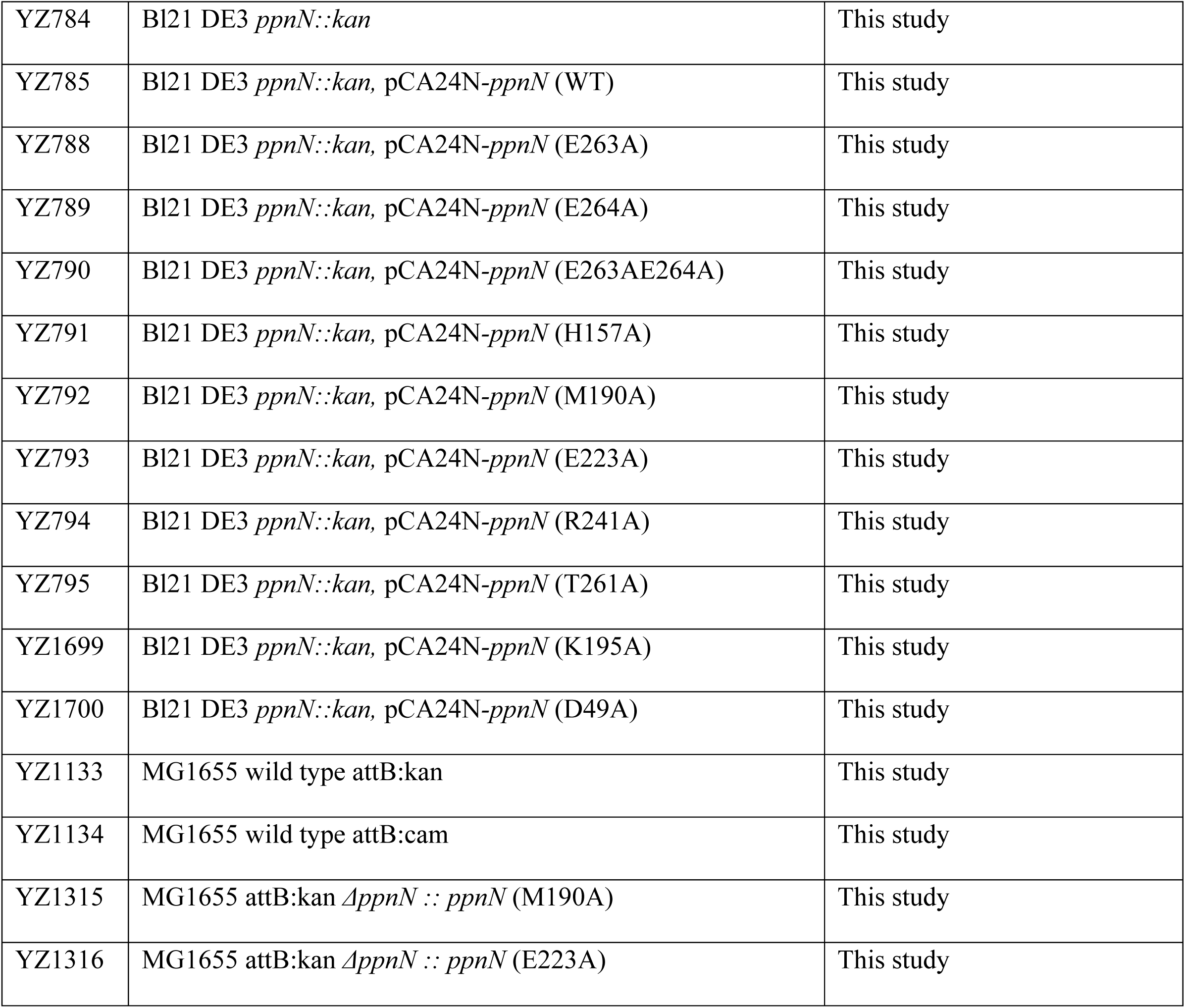
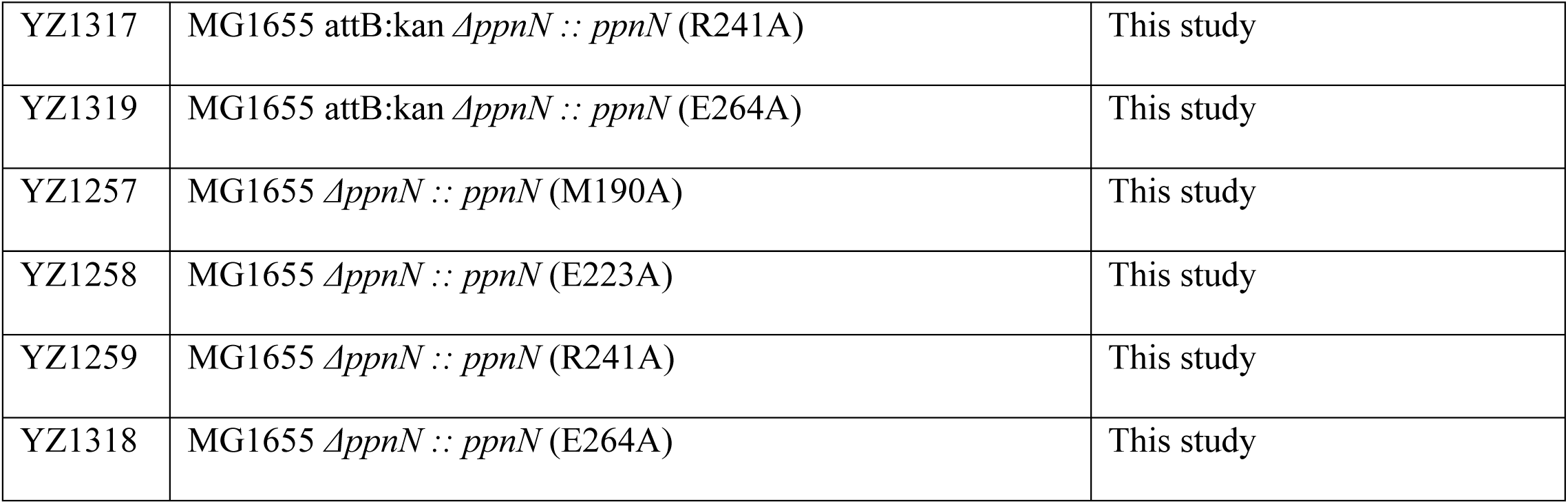
Bacterial strains used in this study.

**Table S3.**
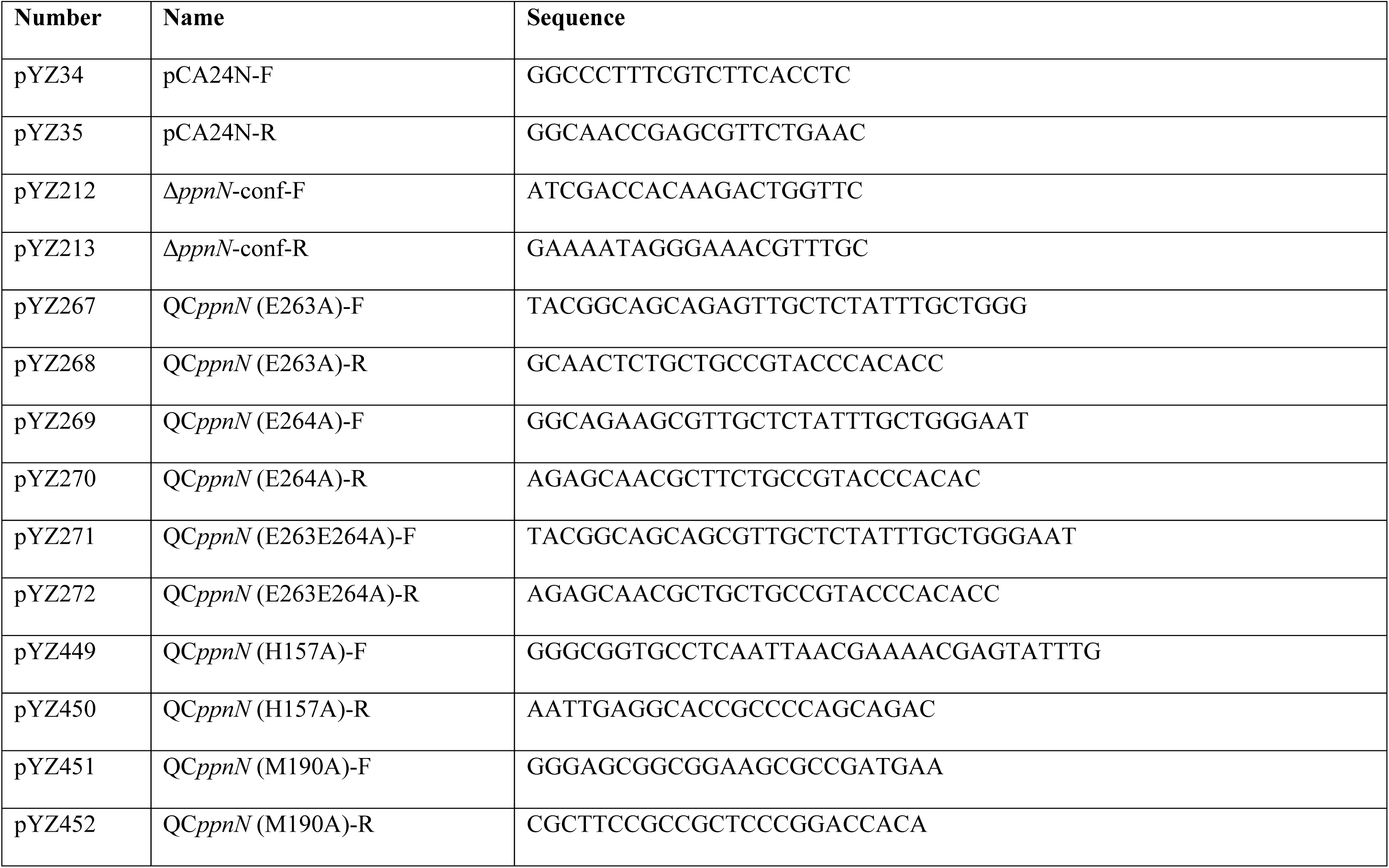
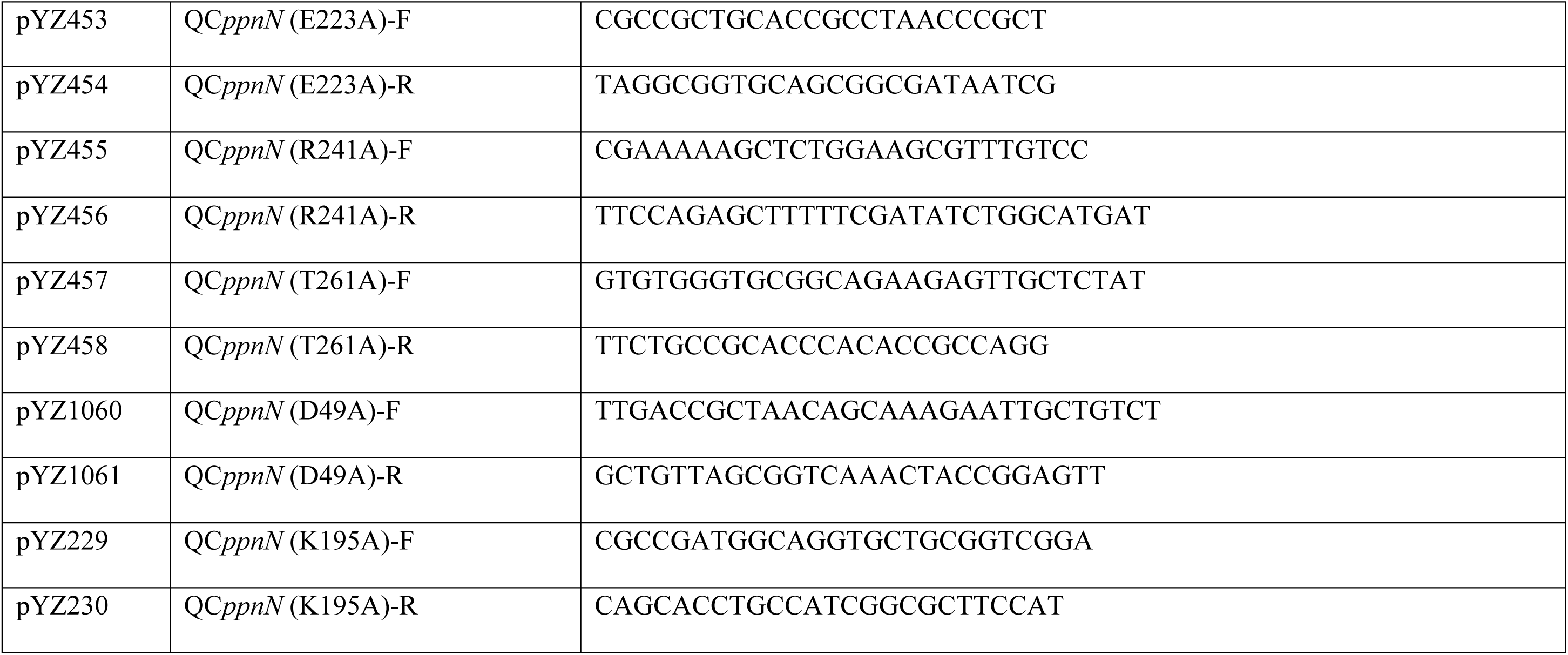
Primers used in this study.

**Table S4.** Descriptor of the binding modes found in the molecular dynamic simulations.

| **MODE #** | **DESCRIPTION** |
| --- | --- |
| **MODE 1** | R341 binds to $\alpha$ and $\epsilon$, R88 binds to $\delta$ |
| **MODE 2** | R341 binds to $\alpha$ and $\epsilon$, R88 binds to $\epsilon$ |
| **MODE 3** | R341 binds to $\beta$, R88 binds to $\epsilon$ |
| **MODE 4** | R341 binds to $\alpha$ and $\delta$, K337 binds to $\epsilon$ |
| **MODE 5** | R341 binds to $\delta$ and $\epsilon$, R88 binds to $\delta$ and K337 binds to $\epsilon$ |
| **MODE 6** | R341 binds to $\epsilon$, R88 binds to $\delta$ |
| **MODE 7** | R88 binds $\epsilon$ |
| **MODE 8** | R341 and R88 binds to $\epsilon$ |
| **MODE 9** | R341 binds to $\alpha$ and $\epsilon$, R88 binds to $\delta$ and $\epsilon$ |
| **MODE 10** | R341 binds to $\delta$ and $\epsilon$, K337 binds to $\epsilon$ |
| **MODE 11** | R341 binds to $\alpha$, R88 binds to $\epsilon$ |
| **MODE 12** | R341 binds to $\alpha$ and $\epsilon$ |
| **MODE 13** | R341 binds to $\alpha$ and $\epsilon$, K337 binds to $\epsilon$ |
| **MODE 14** | R341 binds to $\alpha$, R88 binds to $\delta$, K337 binds to $\epsilon$ |
| **MODE 15** | R341 binds to $\beta$ and $\epsilon$ |
| **MODE 16** | R341 binds to $\alpha$ and $\delta$ |
| **MODE 17** | R341 and K337 binds to $\epsilon$ |
| **MODE 18** | R341 binds to $\delta$, R88 binds to $\epsilon$ |
| **MODE 19** | R341 binds to $\alpha$, R88 binds to $\beta$, K337 binds to $\epsilon$ |
| **MODE 20** | K337 binds to $\epsilon$ |
| **MODE 21** | R341 binds to $\delta$, K337 binds to $\epsilon$ |
| **MODE 22** | R341 binds to $\alpha$, R88 binds to $\delta$ |
| **MODE 23** | R341 binds to $\alpha$, $\delta$ and $\epsilon$, R88 binds to $\epsilon$ |
| **MODE 24** | R341 binds to $\alpha$ and $\delta$, R88 binds to $\epsilon$ |
| **MODE 25** | R341 binds to $\beta$ and $\epsilon$, R88 binds to $\epsilon$ |
| **MODE 26** | R88 binds to $\beta$, K337 binds to $\epsilon$ |
| **MODE 27** | R341 binds to $\epsilon$ |
| **MODE 28** | R341 binds to $\alpha$ and $\epsilon$, R88 binds to $\beta$, K337 binds to $\epsilon$ |
| **MODE 29** | R341 binds to $\alpha$ and $\delta$, R88 binds to $\beta$, K337 binds to $\epsilon$ |
| **MODE 30** | R341 and K337 binds to $\epsilon$, R88 binds to $\beta$ |
| **MODE 31** | R341 binds to $\alpha$, $\beta$ and $\delta$, K337 binds to $\epsilon$ |

**SUPPLEMENTARY Figure 1.**
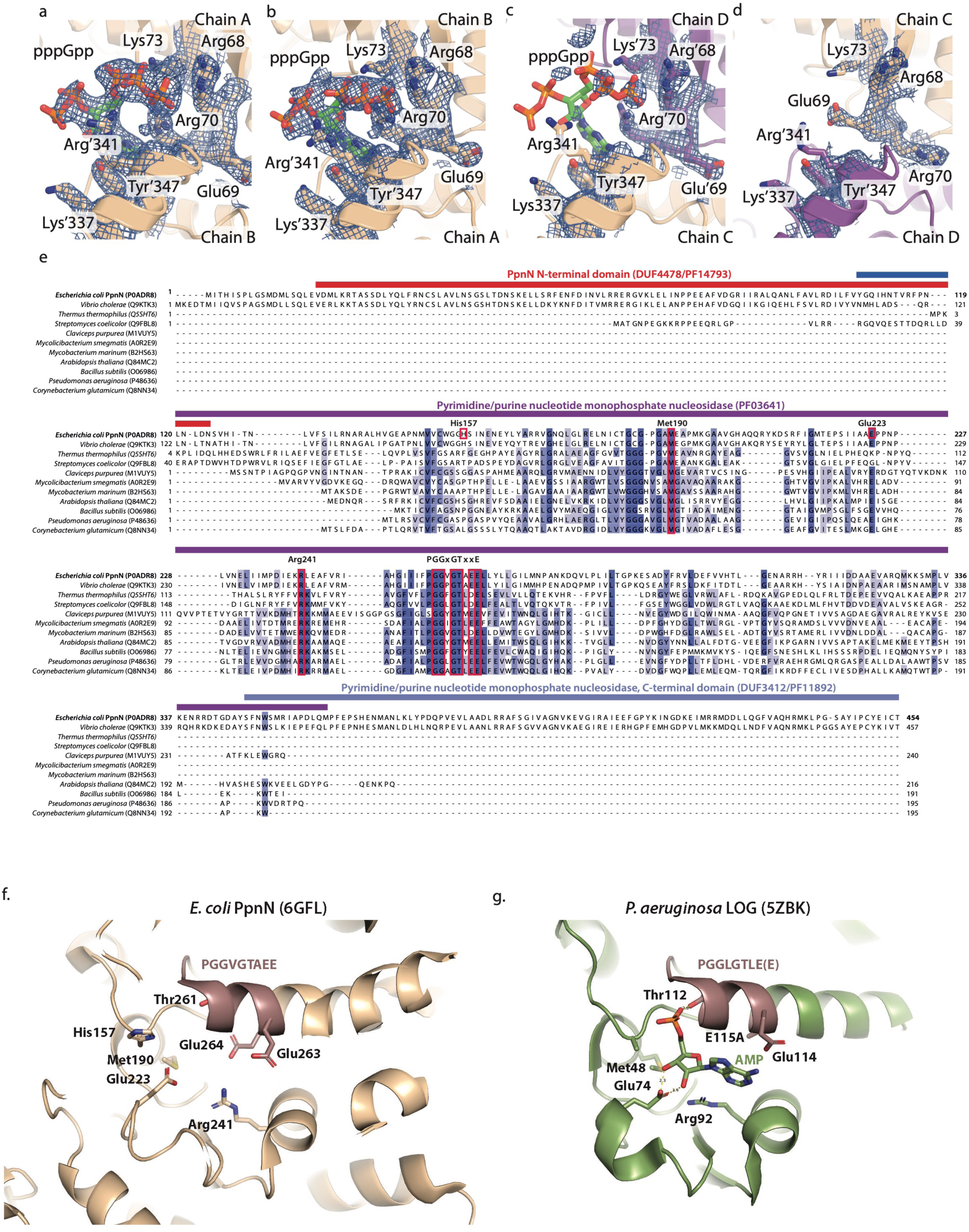
**a**, **b)** Density maps for two alarmone pppGpp molecules in the dimer interfaces between Chain A and Chain B. **c)** The density for the third pppGpp alarmone in the other dimer interface of Chain C and Chain D is much weaker, despite that the coordinating residues are in the same conformation seen in panel-**a,b**. **d)** Density of the fourth, unoccupied alarmone site, where the conformation is in apo state. The 2Fo-Fc maps are set to 1.5 sigma. **e)** Structure-based sequence alignments of homolog proteins of PpnN that have a deposited PDB structure. The PAG1 domain (DUF4478, in red), central catalytic domain (in purple) and PAG2 domain (DUF3412, in blue) are indicated above the sequences. Highly conserved motif PGGxGTxxE and other conserved residues studied are highlighted in red frames. **f, g**) Side-by-side presentation of the three-dimensional poses of key catalytic residues (shown in stick model) of both PpnN and LOG proteins.

**SUPPLEMENTARY Figure 2.**
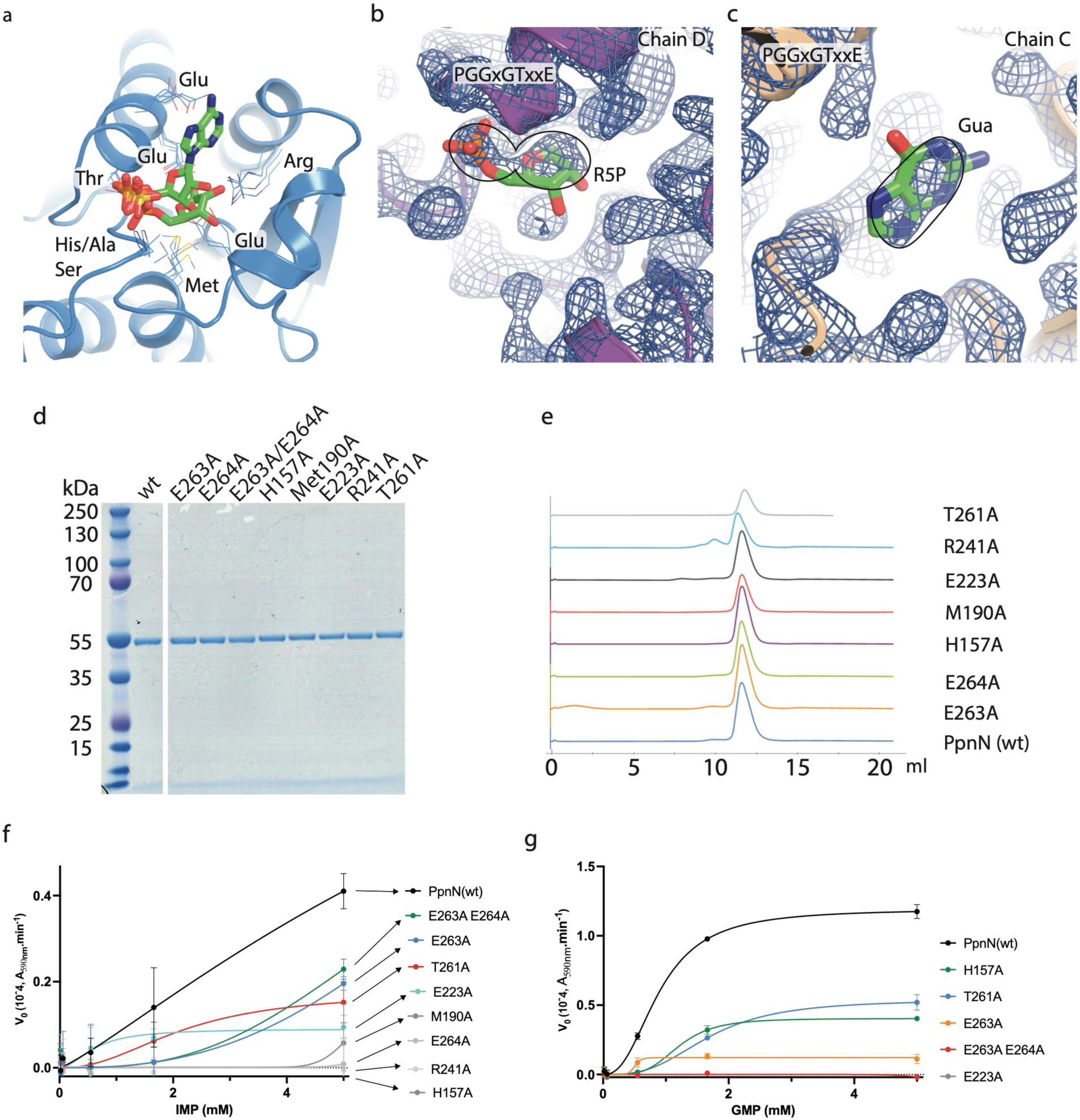
**a)** Conserved active site folds found in homolog LOG proteins out of several organisms. Structures used in the analysis include predicted nucleotide-binding protein from *Idomarina baltica* with SO_4_ ion (PDB ID: 3BQ9^29^), predicted nucleotide-binding protein from *Vibrio cholerae* with PO_4_ ion (PDB ID: 2PMB^28^), uncharacterized protein from *Mycobacterium marinum* with AMP (PDB ID: 3SBX^44^), and phospho-ribo-hydroxylase LOG from *Claviceps purpurea* with R5P (PDB ID: 5AJU^30^). Ligands are shown in stick models, and the coordinating residues are shown as lines. **b)** The density map of the partially open PpnN monomer active site with the modelled R5P; **c)** the density map of the open PpnN monomer active site with the modelled guanine. All 2Fo-Fc maps are set at 1.5 sigma. **d)** SDS-PAGE gel of purified wild type and mutant PpnN proteins (1 μg each). **e)** Size Exclusion Chromatography elution profiles of the wild type and mutant PpnN proteins. **f, g)** Michaelis-Menton enzyme kinetic curves of wt and mutant PpnN proteins by using IMP (**f**) or GMP (**g**) as the substrate. At least three biological replicates were performed, and the curves are fitted using PRISM 10 and an allosteric sigmoidal model. Note the different scales of the Y-axes.

**SUPPLEMENTARY Figure 3.**
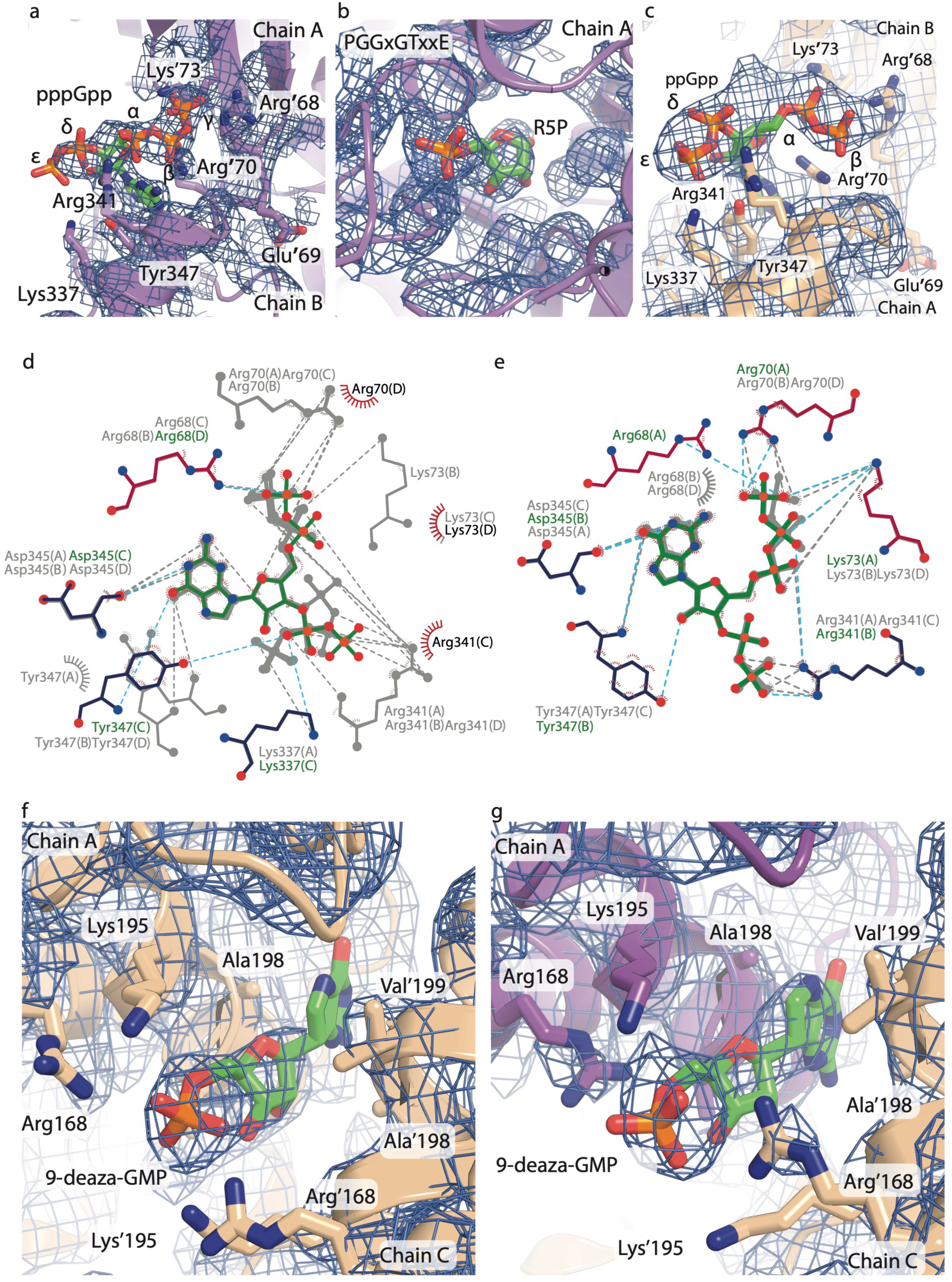
**a)** Density map around pppGpp and **b)** ribose-5’-phosphate (R5P) in the PpnN E264A structure in Figure 3A. The maps are set to 1.5 sigma. **c)** 2Fo-Fc map around ppGpp in the ppGpp-4:4 structure in Figure 3D. **d,e)** LigPlot of ppGpp (d) (from Figure 3D) and pppGpp (e) (from Figure 3A). **f)** One 9-deaza-GMP binding to the dimer interface of two open PpnN monomers and **g)** the other 9- deaza-GMP binding to the dimer interface of one open and one partially open PpnN monomer. The 2Fo-Fc maps are set to 1.5 sigma.

**SUPPLEMENTARY Figure 4.**
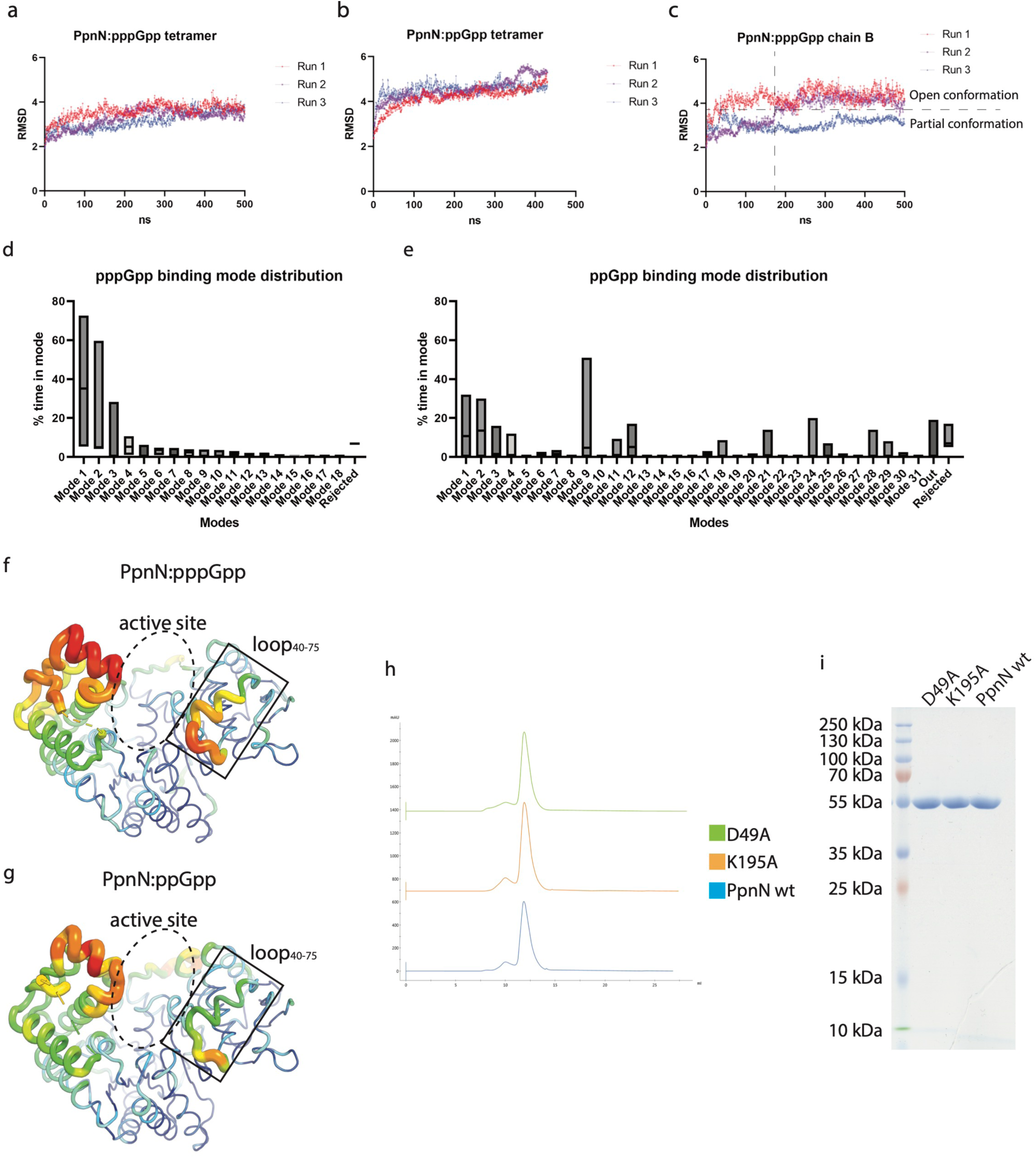
**a-b)** The simulations of both PpnN:pppGpp and PpnN:ppGpp are stable over the course of the simulations. **c)** RMSD plot mirroring that the partially open monomer of PpnN:pppGpp fully opens in the beginning of run 1 (red line), after a while (ca.170 ns) in run 2 (purple line) and remains closed in run 3 (blue line). **d,e)** pppGpp (**d**) adopts fewer, predominant binding modes, whereas **e)** ppGpp adopts more diverse binding modes with less time on each. **f,g)** Temperature factor visualization of PpnN monomers in complex with pppGpp (**f,** from Figure 1A) and with ppGpp (**g,** from Figure 4D). Note the higher temperature -factor of loop_40-75_ in **f** compared to in **g**. Both the loop_40-75_ and the active site are framed up. **h)** Size Exclusion Chromatography elution profiles of the wild type and mutant PpnN proteins. **i)** SDS-PAGE gel of purified wild type and mutant PpnN proteins (1 μg each).

**SUPPLEMENTARY Figure 5.**
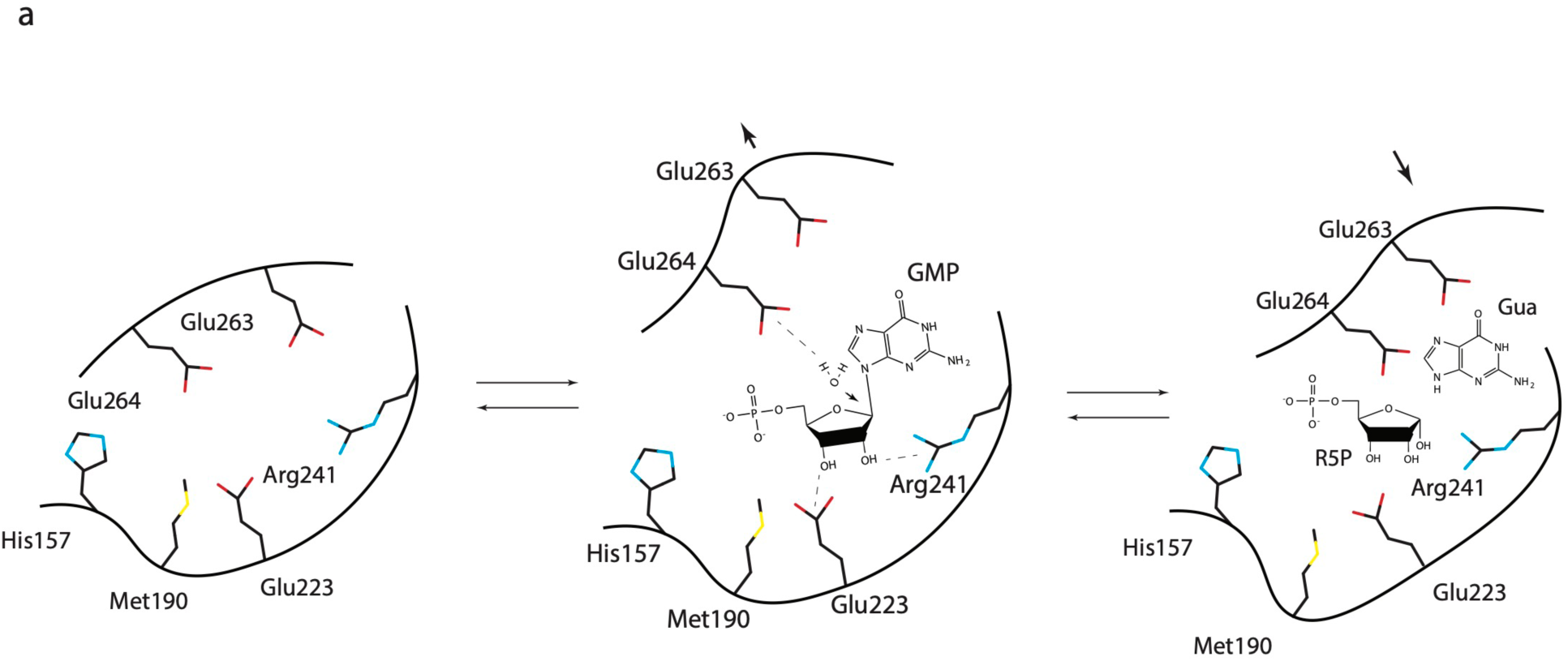
**a)** The proposed catalytic mechanism of PpnN involves the closure of its exposed active site upon binding of the GMP substrate. Key residues, including Glu223 and Arg241, coordinate the 2’ and 3’ hydroxyl groups, while Glu264 coordinates a water molecule priming the N-C bond for hydrolysis. Arrows indicate the movement of key residues during catalysis.

## References

1. Irving, S.E., Choudhury, N.R., and Corrigan, R.M. (2021). The stringent response and physiological roles of (pp)pGpp in bacteria. Nat Rev Microbiol 19, 256–271. 10.1038/s41579-020-00470-y.

2. Wang, B., Dai, P., Ding, D., Del Rosario, A., Grant, R.A., Pentelute, B.L., and Laub, M.T. (2019). Affinity-based capture and identification of protein effectors of the growth regulator ppGpp. Nat Chem Biol 15, 141–150. 10.1038/s41589-018-0183-4.

3. Zhang, Y., Zborníková, E., Rejman, D., and Gerdes, K. (2018). Novel (p)ppGpp Binding and Metabolizing Proteins of Escherichia coli. mBio 9. 10.1128/mBio.02188-17.

4. Durfee, T., Hansen, A.M., Zhi, H., Blattner, F.R., and Jin, D.J. (2008). Transcription profiling of the stringent response in Escherichia coli. J Bacteriol 190, 1084–1096. 10.1128/JB.01092-07.

5. Traxler, M.F., Summers, S.M., Nguyen, H.-T., Zacharia, V.M., Smith, J.T., and Conway, T. (2008). The global, ppGpp-mediated stringent response to amino acid starvation in Escherichia coli. Molecular microbiology 68, 1128–1148. 10.1111/j.1365-2958.2008.06229.x.

6. Sanchez-Vazquez, P., Dewey, C.N., Kitten, N., Ross, W., and Gourse, R.L. (2019). Genome-wide effects on Escherichia coli transcription from ppGpp binding to its two sites on RNA polymerase. Proc Natl Acad Sci U S A. 10.1073/pnas.1819682116.

7. Potrykus, K., Murphy, H., Philippe, N., and Cashel, M. (2011). ppGpp is the major source of growth rate control in E. coli. Environ Microbiol 13, 563–575. 10.1111/j.1462-2920.2010.02357.x.

8. Dalebroux, Z.D., Svensson, S.L., Gaynor, E.C., and Swanson, M.S. (2010). ppGpp conjures bacterial virulence. Microbiol Mol Biol Rev 74, 171–199. 10.1128/MMBR.00046-09.

9. Harms, A., Maisonneuve, E., and Gerdes, K. (2016). Mechanisms of bacterial persistence during stress and antibiotic exposure. Science 354. 10.1126/science.aaf4268.

10. Sevin, D.C., Fuhrer, T., Zamboni, N., and Sauer, U. (2017). Nontargeted in vitro metabolomics for high-throughput identification of novel enzymes in Escherichia coli. Nat Methods 14, 187–194. 10.1038/nmeth.4103.

11. Deo, S.S., Tseng, W.C., Saini, R., Coles, R.S., and Athwal, R.S. (1985). Purification and Characterization of Escherichia-Coli Xanthine-Guanine Phosphoribosyltransferase Produced by Plasmid-Psv2gpt. Biochimica Et Biophysica Acta 839, 233–239. Doi 10.1016/0304-4165(85)90003-0.

12. Guddat, L.W., Vos, S., Martin, J.L., Keough, D.T., and De Jersey, J. (2002). Crystal structures of free, IMP-, and GMP-bound hypoxanthine phosphoribosyltransferase. Protein Sci 11, 1626–1638. 10.1110/ps.0201002.

13. Zhang, Y.E., Baerentsen, R.L., Fuhrer, T., Sauer, U., Gerdes, K., and Brodersen, D.E. (2019). (p)ppGpp Regulates a Bacterial Nucleosidase by an Allosteric Two-Domain Switch. Mol Cell 74, 1239–1249 e1234. 10.1016/j.molcel.2019.03.035.

14. Wang, B., Grant, R.A., and Laub, M.T. (2020). ppGpp Coordinates Nucleotide and Amino-Acid Synthesis in E. coli During Starvation. Mol Cell 80, 29–42.e10. 10.1016/j.molcel.2020.08.005.

15. Grucela, P.K., Fuhrer, T., Sauer, U., Chao, Y., and Zhang, Y.E. (2023). Ribose 5-phosphate: the key metabolite bridging the metabolisms of nucleotides and amino acids during stringent response in Escherichia coli? Microb Cell 10, 141–144. 10.15698/mic2023.07.799.

16. Bærentsen, R.L., Brodersen, D.E., and Zhang, Y.E. (2019). Evolution of the bacterial nucleosidase PpnN and its relation to the stringent response. Microb Cell 6, 450–453. 10.15698/mic2019.09.692.

17. Cortleven, A., Leuendorf, J.E., Frank, M., Pezzetta, D., Bolt, S., and Schmulling, T. (2019). Cytokinin action in response to abiotic and biotic stresses in plants. Plant Cell Environ 42, 998–1018. 10.1111/pce.13494.

18. Cashel, M. (1969). The control of ribonucleic acid synthesis in Escherichia coli. IV. Relevance of unusual phosphorylated compounds from amino acid-starved stringent strains. J Biol Chem 244, 3133–3141.

19. Lazzarini, R.A., Cashel, M., and Gallant, J. (1971). On the regulation of guanosine tetraphosphate levels in stringent and relaxed strains of Escherichia coli. J Biol Chem 246, 4381–4385.

20. Winslow, R.M. (1971). A consequence of the rel gene during a glucose to lactate downshift in Escherichia coli. The rates of ribonucleic acid synthesis. J Biol Chem 246, 4872–4877.

21. Mechold, U., Potrykus, K., Murphy, H., Murakami, K.S., and Cashel, M. (2013). Differential regulation by ppGpp versus pppGpp in Escherichia coli. Nucleic Acids Res 41, 6175–6189. 10.1093/nar/gkt302.

22. Keasling, J.D., Bertsch, L., and Kornberg, A. (1993). Guanosine pentaphosphate phosphohydrolase of Escherichia coli is a long-chain exopolyphosphatase. Proc Natl Acad Sci U S A 90, 7029–7033.

23. Kihira, K., Shimizu, Y., Shomura, Y., Shibata, N., Kitamura, M., Nakagawa, A., Ueda, T., Ochi, K., and Higuchi, Y. (2012). Crystal structure analysis of the translation factor RF3 (release factor 3). FEBS Lett 586, 3705–3709. 10.1016/j.febslet.2012.08.029.

24. Hwang, J., and Inouye, M. (2008). RelA functionally suppresses the growth defect caused by a mutation in the G domain of the essential Der protein. J Bacteriol 190, 3236–3243. 10.1128/jb.01758-07.

25. Steinchen, W., Schuhmacher, J.S., Altegoer, F., Fage, C.D., Srinivasan, V., Linne, U., Marahiel, M.A., and Bange, G. (2015). Catalytic mechanism and allosteric regulation of an oligomeric (p)ppGpp synthetase by an alarmone. Proc Natl Acad Sci U S A 112, 13348–13353. 10.1073/pnas.1505271112.

26. Naseem, M., Bencurova, E., and Dandekar, T. (2018). The Cytokinin-Activating LOG-Family Proteins Are Not Lysine Decarboxylases. Trends Biochem Sci 43, 232–236. 10.1016/j.tibs.2018.01.002.

27. Naseem, M., Sarukhanyan, E., and Dandekar, T. (2015). LONELY-GUY Knocks Every Door: Crosskingdom Microbial Pathogenesis. Trends Plant Sci 20, 781–783. 10.1016/j.tplants.2015.10.017.

28. Patskovsky, Y., Toro, R., Meyer, A.J., Dickey, M., Eberle, M., Koss, J., Groshong, C., Wasserman, S.R., Sauder, J.M., Burley, S.K., Almo, S.C. (2007). Crystal structure of predicted nucleotide-binding protein from Vibrio cholerae. N.Y.S.R.C.f.S.G. (NYSGXRC), ed.

29. Patskovsky, Y., Toro, R., Meyer, A.J., Dickey, M., Eberle, M., Koss, J., Groshong, C., Wasserman, S.R., Sauder, J.M., Burley, S.K., Almo, S.C. (2008). Crystal structure of predicted nucleotide-binding protein from Idiomarina baltica OS145. N.Y.S.R.C.f.S.G. (NYSGXRC), ed.

30. Dzurová, L., Forneris, F., Savino, S., Galuszka, P., Vrabka, J., and Frébort, I. (2015). The three-dimensional structure of "Lonely Guy" from provides insights into the phosphoribohydrolase function of Rossmann fold-containing lysine decarboxylase-like proteins. Proteins 83, 1539–1546. 10.1002/prot.24835.

31. Jeon, W.B., Allard, S.T.M., Bingman, C.A., Bitto, E., Han, B.W., Wesenberg, G.E., and Phillips, G.N. (2006). X-ray crystal structures of the conserved hypothetical proteins from gene loci At5g11950 and At2g37210. Proteins 65, 1051–1054. 10.1002/prot.21166.

32. Seo, H., and Kim, K.J. (2018). Structural insight into molecular mechanism of cytokinin activating protein from Pseudomonas aeruginosa PAO1. Environ Microbiol 20, 3214–3223. 10.1111/1462-2920.14287.

33. Helenius, M., Jalkanen, S., and Yegutkin, G. (2012). Enzyme-coupled assays for simultaneous detection of nanomolar ATP, ADP, AMP, adenosine, inosine and pyrophosphate concentrations in extracellular fluids. Biochim Biophys Acta 1823, 1967–1975. 10.1016/j.bbamcr.2012.08.001.

34. Kronborg, K., and Zhang, Y. (2023). Substrate promiscuity of the Escherichia coli xanthine oxidase. In U.o. Copenhagen, ed. bioRxiv.

35. Blank, K., Hensel, M., and Gerlach, R.G. (2011). Rapid and highly efficient method for scarless mutagenesis within the Salmonella enterica chromosome. PLoS One 6, e15763. 10.1371/journal.pone.0015763.

36. Rymer, R.U., Solorio, F.A., Tehranchi, A.K., Chu, C., Corn, J.E., Keck, J.L., Wang, J.D., and Berger, J.M. (2012). Binding Mechanism of Metal•NTP Substrates and Stringent-Response Alarmones to Bacterial DnaG-Type Primases. Structure 20, 1478–1489. 10.1016/j.str.2012.05.017.

37. Mechold, U., Potrykus, K., Murphy, H., Murakami, K.S., and Cashel, M. (2013). Differential regulation by ppGpp versus pppGpp in. Nucleic Acids Research 41, 6175–6189. 10.1093/nar/gkt302.

38. Pausch, P., Steinchen, W., Wieland, M., Klaus, T., Freibert, S.A., Altegoer, F., Wilson, D.N., and Bange, G. (2018). Structural basis for (p)ppGpp-mediated inhibition of the GTPase RbgA. Journal of Biological Chemistry 293, 19699–19709. 10.1074/jbc.RA118.003070.

39. Czech, L., Mais, C.N., Kratzat, H., Sarmah, P., Giammarinaro, P., Freibert, S.A., Esser, H.F., Musial, J., Berninghausen, O., Steinchen, W., et al. (2022). Inhibition of SRP-dependent protein secretion by the bacterial alarmone (p)ppGpp. Nat Commun 13. 10.1038/s41467-022-28675-0.

40. Chau, N.Y.E., Perez-Morales, D., Elhenawy, W., Bustamante, V.H., Zhang, Y.E., and Coombes, B.K. (2021). (p)ppGpp-Dependent Regulation of the Nucleotide Hydrolase PpnN Confers Complement Resistance in Salmonella enterica Serovar Typhimurium. Infect Immun 89. 10.1128/IAI.00639-20.

41. Launay, A. (2016). Study of the emergence of the diversity of Escherichia coli in vivo by whole genome sequencing. Universite ’ Pierre et Marie Curie - Paris VI.

42. Klemm, E.J., Gkrania-Klotsas, E., Hadfield, J., Forbester, J.L., Harris, S.R., Hale, C., Heath, J.N., Wileman, T., Clare, S., Kane, L., et al. (2016). Emergence of host-adapted Salmonella Enteritidis through rapid evolution in an immunocompromised host. Nature Microbiology 1, 15023–15023. 10.1038/nmicrobiol.2015.23.

43. Samanovic, M.I., Tu, S., Novák, O., Iyer, L.M., McAllister, F.E., Aravind, L., Gygi, S.P., Hubbard, S.R., Strnad, M., and Darwin, K.H. (2015). Proteasomal control of cytokinin synthesis protects Mycobacterium tuberculosis against nitric oxide. Mol Cell 57, 984–994. 10.1016/j.molcel.2015.01.024.

44. Baugh, L., Phan, I., Begley, D.W., Clifton, M.C., Armour, B., Dranow, D.M., Taylor, B.M., Muruthi, M.M., Abendroth, J., Fairman, J.W., et al. (2015). Increasing the structural coverage of tuberculosis drug targets. Tuberculosis (Edinb) 95, 142–148. 10.1016/j.tube.2014.12.003.

45. Winn, M.D., Ballard, C.C., Cowtan, K.D., Dodson, E.J., Emsley, P., Evans, P.R., Keegan, R.M., Krissinel, E.B., Leslie, A.G.W., McCoy, A., et al. (2011). Overview of the CCP4 suite and current developments. Acta Crystallogr D 67, 235–242. 10.1107/S0907444910045749.

46. Mccoy, A.J., Grosse-Kunstleve, R.W., Adams, P.D., Winn, M.D., Storoni, L.C., and Read, R.J. (2007). Phaser crystallographic software. J Appl Crystallogr 40, 658–674. 10.1107/S0021889807021206.

47. Smart, O.S., Womack, T.O., Flensburg, C., Keller, P., Paciorek, W., Sharff, A., Vonrhein, C., and Bricogne, G. (2012). Exploiting structure similarity in refinement: automated NCS and target-structure restraints in. Acta Crystallogr D 68, 368–380. 10.1107/S0907444911056058.

48. Afonine, P.V., Grosse-Kunstleve, R.W., Echols, N., Headd, J.J., Moriarty, N.W., Mustyakimov, M., Terwilliger, T.C., Urzhumtsev, A., Zwart, P.H., and Adams, P.D. (2012). Towards automated crystallographic structure refinement with phenix.refine. Acta Crystallogr D 68, 352–367. 10.1107/S0907444912001308.

49. Emsley, P., Lohkamp, B., Scott, W.G., and Cowtan, K. (2010). Features and development of Coot. Acta Crystallographica Section D-Biological Crystallography 66, 486–501. 10.1107/S0907444910007493.

50. Murshudov, G.N., Skubák, P., Lebedev, A.A., Pannu, N.S., Steiner, R.A., Nicholls, R.A., Winn, M.D., Long, F., and Vagin, A.A. (2011). REFMAC5 for the refinement of macromolecular crystal structures. Acta Crystallogr D 67, 355–367. 10.1107/S0907444911001314.

51. Emsley, P. (2017). Tools for ligand validation in Coot. Acta Crystallogr D 73, 203–210. 10.1107/S2059798317003382.

52. Schüttelkopf, A.W., and van Aalten, D.M.F. (2004). PRODRG: a tool for high-throughput crystallography of protein-ligand complexes. Acta Crystallogr D 60, 1355–1363. 10.1107/S0907444904011679.

53. Liebschner, D., Afonine, P.V., Moriarty, N.W., Poon, B.K., Sobolev, O.V., Terwilliger, T.C., and Adams, P.D. (2017). Polder maps: improving OMIT maps by excluding bulk solvent. Acta Crystallogr D 73, 148–157. 10.1107/S2059798316018210.

54. Van der Spoel, D., Lindahl, E., Hess, B., Groenhof, G., Mark, A.E., and Berendsen, H.J.C. (2005). GROMACS: Fast, flexible, and free. J Comput Chem 26, 1701–1718. 10.1002/jcc.20291.

55. Bas, D.C., Rogers, D.M., and Jensen, J.H. (2008). Very fast prediction and rationalization of pKa values for protein-ligand complexes. Proteins 73, 765–783. 10.1002/prot.22102.

56. Huang, J., and MacKerell, A.D. (2013). CHARMM36 all-atom additive protein force field: Validation based on comparison to NMR data. J Comput Chem 34, 2135–2145. 10.1002/jcc.23354.

57. Steinchen, W., and Bange, G. (2016). The magic dance of the alarmones (p)ppGpp. Mol Microbiol 101, 531–544. 10.1111/mmi.13412.

58. Kushwaha, G.S., Patra, A., and Bhavesh, N.S. (2020). Structural Analysis of (p)ppGpp Reveals Its Versatile Binding Pattern for Diverse Types of Target Proteins. Frontiers in Microbiology 11. 10.3389/fmicb.2020.575041.

59. Humphrey, W., Dalke, A., and Schulten, K. (1996). VMD: Visual molecular dynamics. J Mol Graph Model 14, 33–38. Doi 10.1016/0263-7855(96)00018-5.

60. Kitagawa, M., Ara, T., Arifuzzaman, M., Ioka-Nakamichi, T., Inamoto, E., Toyonaga, H., and Mori, H. (2006). Complete set of ORF clones of Escherichia coli ASKA library (A Complete Set of E. coli K-12 ORF Archive): Unique Resources for Biological Research. DNA Research 12, 291–299. 10.1093/dnares/dsi012.

